# Maternal Thyroid Supplementation Prevents Autistic-relevant Social Behavior and Hypothalamic Oxytocin Depletion Produced by Developmental Exposure to Environmental Toxicants

**DOI:** 10.1101/2025.07.28.667293

**Authors:** Elena V. Kozlova, Maximillian E. Denys, Anthony E. Bishay, Crystal N. Luna, Meri De Angelis, Luis Campoy, Amna Habbal, Artha A. Lam, Naran Luvsanravdan, Anastasia Ghilenschi Colton, David Carter, Timo Müller, Karl-Werner Schramm, Margarita C. Currás-Collazo

## Abstract

Environmental toxicants that target the developing brain are suspected of contributing to autism spectrum disorder risk but causative evidence is lacking. We and others have shown that the indoor flame retardants, polybrominated diphenyl ethers (PBDEs), reduce prosocial behavior, however, few studies have assessed the central targets and underlying mechanisms. PBDEs are well established endocrine disruptors of the expanded thyroid system, which also regulates the prosocial neuropeptides oxytocin (OXT) and vasopressin (AVP) and their hypothalamic signaling. The potential role of PBDE-induced thyroid hormone (TH) deregulation in mediating disruption of central OXT and ASD-like social behavior deficits remains unmapped. To address this gap, we conducted a study in C57BL6/N mice that examined behavioral and neuromolecular reprogramming after developmental exposure to the commercial PBDE mixture, DE-71, and evaluated the therapeutic potential of TH supplementation. Dams were exposed daily during gestation and lactation to corn oil vehicle, low dose (0.1 mg/kg) and high dose (0.4 mg/kg) of DE-71 with or without concurrent L-thyroxine (+mT4). In offspring, dose-dependent ASD-relevant behavioral responses and central neuroendocrine OXT neuron depletion after developmental PBDE exposure was prevented with mT4. mRNA transcripts for the TH transporter *Mct8*, deiodinase (*Dio3)* and estrogen receptor beta (*Esr2*) expressed on OXT neurons in PVH were upregulated in low dose females. In contrast, *Mct8* and *Dio3* were downregulated in low dose males. These findings uncover sex-specific mechanisms of PBDE-induced reprogramming of TH-regulated pathways in hypothalamic neuroendocrine cells leading to depleted central OXT signaling and ultimately ASD-relevant phenotypes. Importantly, we provide novel evidence of the therapeutic potential of maternal thyroid supplementation against toxicant-induced neurodevelopmental disorders.

## Introduction

Autism Spectrum Disorder (ASD) is a neurodevelopmental condition characterized by alterations across two core symptom domains: social communication and interaction; and restricted, repetitive patterns of behavior, interests, or activities [1, 2]. The prevalence of ASD has increased significantly to 1 in 31 of U.S. children, representing a nearly 5-fold increase since 2000 [3]. ASD affects each sex differently and is 3.4 times more prevalent in boys [3]. Factors such as increased awareness, improved detection and broadening of diagnostic criteria contribute to but cannot fully account for this epidemic rise [4]. The etiology of ASD is not well understood but both genetic vulnerability [5] and environmental triggers [6], are likely to confer ASD risk in susceptible individuals through gene x environment interactions (G x E) [7]. The heterogeneous and multifactorial nature of ASD hinders the identification of reliable biomarkers and no pharmacological treatments are available that effectively target its core symptom domains [2].

Polybrominated Diphenyl Ethers (PBDEs) are one candidate environmental factor that are suspected of conferring ASD risk. PBDEs are a class of persistent organic pollutants (POPs) that have been added as flame retardants to plastics, textiles and electronics for decades [8]. As superficial additives, PBDEs leach, volatilize, and photodegrade over time; and due to their lipophilicity and long half-lives, bioaccumulate in biota, including maternal placenta, serum, breast milk and fetal tissues [9–12]. Out of growing concern for their developmental neurotoxicity [8], PBDEs were banned and/or voluntary phased-out in the US. As a result, most biomonitoring studies around the world have observed slowly declining trends in PBDE levels [13–15]. However, the rate of decline in biota varies inconsistently across different congeners, age, and gender [16–18]. Thus, despite the phase out, PBDE congeners are still widely detected in humans [19].

Moreover, PBDE contamination is ongoing due to environmental reintroduction through e-waste, recycled microplastic vectors, and recycling that is estimated to continue through 2050 [20, 21].

Evidence from epidemiological studies supports an association between PBDE exposure and characteristics of neurodevelopmental disorders (NDDs) such as impairments in executive function, poor attention and behavioral regulation, reduced social scores, and lower IQ in children [22–26], while other findings are less clear [27–29]. Notably, Hamra et al. (2021) found that each one-unit increase in the weighted sum of PBDEs was associated with a 1.41-fold increase in the odds of ASD, as measured by the Social Responsiveness Scale (SRS).[30]. However, critical scientific, regulatory, and economic barriers still exist to causally link specific environmental exposures to specific psychosocial NDDs [31].

Murine studies examining the effects of brominated frame retardants (BFR) report disrupted social cognition across several domains [32–34], while other studies found null effects in the absence of a second environmental hit, possibly due to heterogeneity of BFR type, timing of exposure, sex and/or model organism [33–36]. Compounding this heterogeneity, PBDE congeners show estrogenic, anti-estrogenic and/or androgenic actions [37] that may produce sex-specific effects. Recently, we have shown that PBDE-exposed F1 female progeny display autism-relevant characteristics such as deficient social recognition memory (SRM), exaggerated repetitive behavior, deficient social odor discrimination, and deregulated markers for prosocial hypothalamic genes [38]. The potential mechanisms underlying these abnormal and persistent phenotypes are unknown but may be related to the thyroid-disrupting actions of PBDEs and interference with the TH-mediated regulation of prosocial neuroendocrine systems [39].

Recent epidemiological studies have demonstrated that alterations to thyroid hormone (TH) function during pregnancy may be a contributing factor to ASD etiology [40–44]. However, while insufficient TH levels during pregnancy can lead to long-term intellectual and behavioral impairments in children, the importance of TH insufficiency in ASD has not been established [45]. Clinically, thyroid supplementation in mothers or children has been used to treat congenital hypothyroidism but not other NDDs such as ASD [46]. The main TH, thyroxine (T4), and its active deiodinated form, triiodothyronine (T3) are essential epigenetic factors that have a permissive role in regulating the precise spatiotemporal development of the brain. These include the assembly of functional neuronal circuits through the TH-mediated progressive events of neurogenesis, cell migration, synaptogenesis, formation of cortical layers, neuronal and glial differentiation and myelination [47, 48]. Given the role of THs in neurodevelopment, understanding how toxicant-induced TH disruption influences ASD risk is critical for informing early interventions and prevention strategies.

Human and animal studies have identified that the main endocrine disrupting effects of PBDEs are associated with endocrine disruption of thyroid and reproductive hormones [49–55]. However, few studies have examined disruption of other endocrine axes (see reviews [39, 56, 57]). Oxytocin (OXT) and vasopressin (AVP), nonapeptide hormones synthesized in the neuroendocrine hypothalamus [58], are critical for social recognition ability and have been implicated in both the pathogenesis [59, 60] and treatment [61–64] of psychosocial NDDs such as ASD. OXT/AVP systems are part of a multi-node brain network that regulates several social domains including social recognition, social approach, pair bonding, maternal behavior, reward and social and emotional discrimination [65–68]. In humans, signaling within this social neural network (SNN) is altered in autistic individuals [69]. Interestingly, several animal studies have shown that THs regulate hypothalamic OXT/AVP mRNA and content [70, 71], and axonal release [70–73]. Gene transcription of OXT mRNA in both cellular models and living animal brains is regulated by THs transported across plasma membranes and into the nucleus where the T3-activated receptor binds to TH response element (TRE) on the OXT promoter. Similarly, the *Oxt* gene contains an estrogen response element (ERE) that enables estrogen receptor β (ERβ) to modulate *Oxt* expression [74]. Taken together, this evidence supports the premise that central TH actions may contribute to social behavior by regulating social neuropeptides and the structural development of the SNN.

We and others have also shown that the paraventricular (PVH) and supraoptic (SON) nuclei of the hypothalamus, the main sites of OXT/AVP synthesis, are vulnerable to endocrine disruption by PBDEs and other EDCs [38, 75–78]. PBDEs may impair the central release of oxytocin (OXT) from somatodendritic compartments [79] and axonal projections of OXT-ergic neurons in the paraventricular nuclei (PVH) of the hypothalamus targeting SNN nuclei [80], potentially contributing to the risk of NDDs. Supporting the somatodendritic effects, PBDEs, and structurally related organochlorines found in Aroclor 1254, reduce AVP release from the somatodendritic compartments of magnocellular neuroendocrine cells located in the SON [75, 76].

Because PBDEs are prominent thyroid-disrupting chemicals, in the current study, we hypothesized that developmental DE-71 effects on social behavior are coincident with brain OXT and TH status disruption. We also tested the novel hypothesis that the abnormal phenotypes induced by DE-71 can be prevented through maternal thyroid hormone supplementation, in order to determine whether ASD-relevant effects occur in a thyroid-dependent manner.Our results show that perinatal exposure to PBDEs alters offspring brain TH status in an age- and sex-specific manner and deplete PVH and SON hypothalamic OXT, effects that correlate with ASD-like behavior. In addition, female and male offspring displayed differential changes in mRNA markers of TH transport and deiodination and estrogen receptors. Maternal T4 supplementation prevented many of these effects highlighting potential biomarkers and mechanisms contributing to ASD in offspring. These findings may have implications for ascribing risk of neurodevelopmental disorders to environmental chemicals and for clinical application of maternal thyroid supplementation.

## Methods

### Animal Care and Maintenance

C57BL/6N mice were obtained from Charles River Laboratories (Raleigh, NC, USA). Animals were housed in a controlled environment: temperature (21.1–22.8 °C), light (12:12 light– dark; lights on at 0800 h), and humidity (20-70%). Mice had access to food and water *ad libitum*. Care and treatment of animals was performed in compliance with guidelines from NIH and approved by the University of California Riverside Institutional Animal Care and Use Committee (AUP# 5, 20170026 and 20200018). The studies were carried out in accordance with the ARRIVE 2.0 guidelines.

### Maternal PBDE Exposure and TH Manipulation

As described previously, C57BL6/N mouse dams (F0) were exposed to the commercial penta-BDE mixture, DE-71, via oral cornflake treats at 0.1 (L-DE-71) or 0.4 mg/kg bw/d (H-DE-71) from 3 wk prior to gestation (GD) through end of lactation on postnatal day (P)21 as described [81][38]. DE-71 dosing solutions were prepared in corn oil vehicle (VEH/CON) using 2 mL of stock solution/kg bw. Controls received vehicle (VEH/CON). Maternal thyroxine supplementation (mT4) was achieved by treating VEH/CON and DE-71 treated dams with L-thyroxine sodium salt pentahydrate (T4; TCI America, USA, T0245) via drinking water from GD12 through P21 as described [82]. In brief, a stock T4 solution (250 μg/mL) containing bovine serum albumin (0.01%) was prepared weekly and used to make a diluted working T4 solution (0.25 ug/mL) using tap water. Maternal T4 administration was based on an average dam water intake of 7.4 mL/d (GD15–P0) and 13.33 mL/d (P0–P10), resulting in estimated T4 doses of 1.9 µg/d and 3.3 µg/d, corresponding to approximately 5.3 µg/100 g bw and 9.5 µg/100 g bw, respectively. A separate group of VEH/CON dams were administered 50 mg/L (50 ppm) propylthiouracil (PTU, Frontier Scientific Inc., USA, 254393) dissolved in 0.1% DMSO via drinking water from GD14-P21 [83]. Dam food and water intake, weight gain and litter size and sex ratio were monitored (**Supplementary Table 1**). At P0, a subset of dams were subjected to tail bleed for plasma total T4 (TT4), free T4 and glucose measurement. At P3, a pup retrieval test was performed. Plasma collected at sacrifice (P22) was used to measure plasma T4 and OXT. Offspring were tested in the postnatal, juvenile or adult periods (**Fig. 1**) Cohort 1 offspring were used for analysis of plasma TT4, brain TH species and gene expression. Cohort 2 offspring were perfused at P30-40 and brains processed for immunohistochemistry, RNA *in situ* hybridization and blood draw at sacrifice for OXT measurements. Cohort 3 offspring underwent behavioral testing. Every experiment tested subjects from at least 3 litters per group. Biological replicates are specified in figure legends.

**Figure 1.**
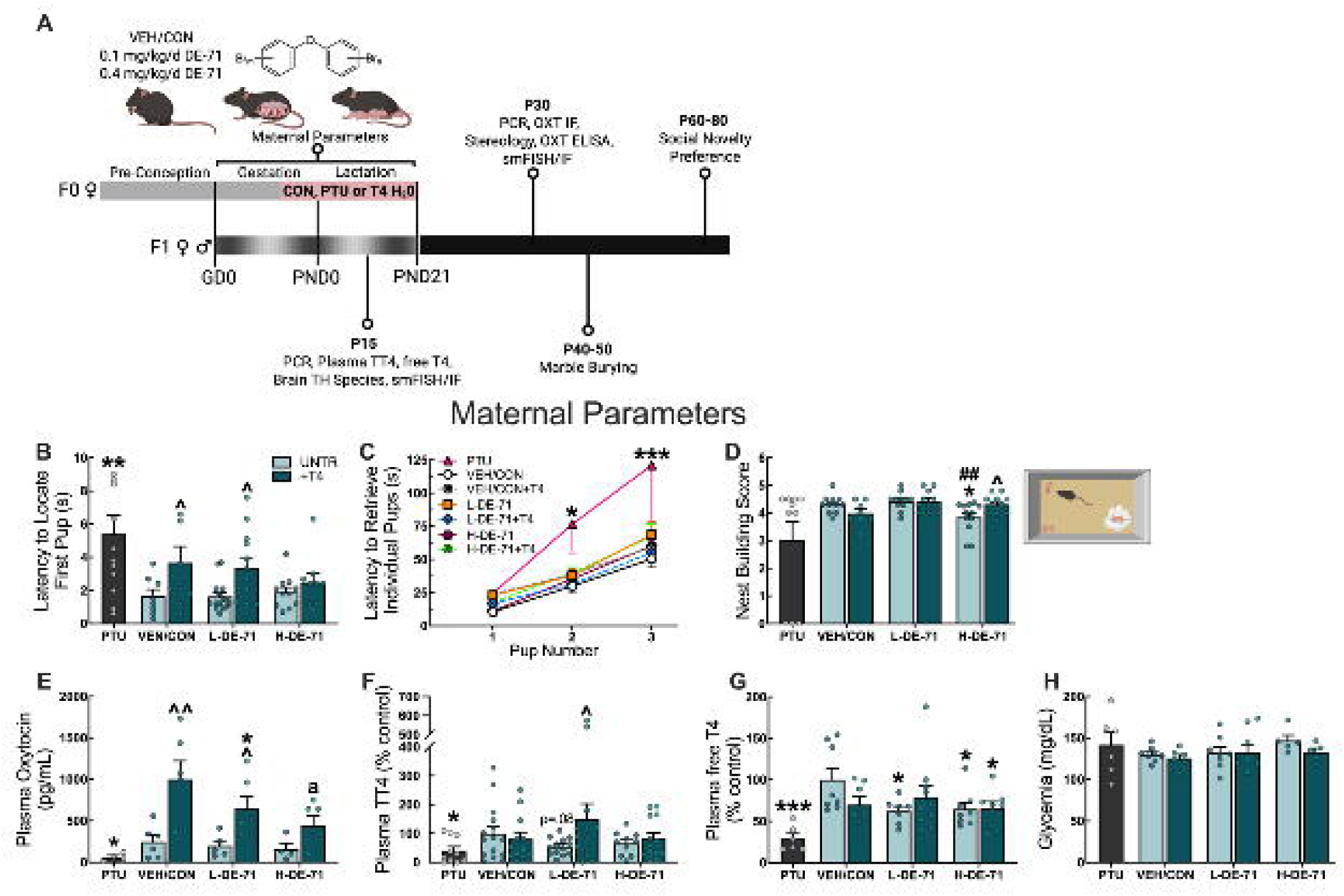
Experimental design and maternal parameters measured. (A) Timeline of DE-71 exposure in dams (F0♀) and offspring (F1) and *in vivo* and *ex vivo* testing. Maternal parameters including weight gain, food and water intake (GD1 to P10), nest scores, pup retrieval, and plasma THs, OXT and glucose were measured GD0-P22. Pups were exposed to DE-71 indirectly during development (hatched shading) for ∼39 d (GD0-P21). A subgroup of VEH/CON and DE-71-exposed dams received maternal levothyroxine supplementation (T4) at GD12-P21 and another received only the anti-thyroid drug, PTU, at GD14-P21 via drinking water. (B, C) Pup retrieval by dam mothers was measured at P2-8 as latency to *locate* the first pup (B) and latency to *retrieve* pups back to nest (C). (D) Nest building scores during P2-6. (E) Dam plasma OXT at P22. (F) Dam plasmaTT4 at P0. (G) Dam plasma free T4 at P0. (H) Dam glycemia at P0. *statistical difference vs VEH/CON (\**p*<0.05, \*\**p*<0.01, \*\*\**p*<0.001). ^statistical difference vs corresponding untreated group (^*p*<0.05, ^^*p*<0.01). ^##^statistical difference vs L-DE-71, *p*<0.01). ^a^statistical difference vs VEH/CON+mT4, *p*<0.05. Sample size (*n*): 8-15/group (B-D); 4-15/group (E-H). GD, gestational day; IF: immunofluorescence; T4: maternal levothyroxine supplementation; mPTU: maternal treatment with propylthiouracil; OXT: oxytocin; P: postnatal day; smFISH: single-molecule *in situ* hybridization; TT4: total T4

### Maternal Nest Scores

The sophistication of nests built by dams from two pressed cotton nestlets (5 × 5 cm) were evaluated at PND 0–8 using as described [38]. Bland-Altman plots indicate agreement and negligible skewing in scores provided by 2 observers (**Supplementary** Fig. 1).

### Liquid Chromatography–Mass Spectrometry (LC/MS) for Brain TH Determination

Mice were sacrificed under isoflurane anesthesia followed by cervical dislocation. Whole brains were rapidly dissected and flash-frozen. Samples were homogenized at -20°C to powder from which ∼100 mg was extracted for determination of THs. The standard solutions and clean-up procedure was performed as previously described [84]. The detection and quantification of seven THs: L-Thyroxine (T4), 3,3′,5-triiodothyronine (T3), reverse 3,3’,5’-triiodothyronine (rT3), 3,3′-diiodo-L-thyronine (T2), 3,5-diiodo-L-thyronine (rT2), 3-iodo-L-thyronine (T1) and 3-iodothyronamine (T1AM) was performed with Sciex QTrap 5500 mass spectrometer interfaced with Agilent 1290 Infinity II LC system or with Shimadzu Nexera X2 LC system. For both configurations, the flow rate was set at 0.3 mL/min, the column used was a Zorbax Eclipse Plus C18 (2.1x50 mm, 1.8 uM) with a column temperature fixed at 40°C. The injection volume was always 10 μL. The mobile phases were water (A) and acetonitrile (B), each containing 0.1% formic acid (v/v). Gradient elution was the same for both instrumentations and is reported in **Supplementary Table 2**. Analytes were detected using an electrospray ionization source (ESI) in positive mode. The tandem mass spectrometer was operated under multiple reaction monitoring mode (MRM). The MS/MS parameters for both configurations were the same and indicated in **Supplementary Table 3**. Data acquisition, linearity of the standard curves and quantification of the samples were performed using Analyst software. rT2, T1, T1AM were below the method detection limit (MDL).

Blood was centrifuged at 2000 × g at 4°C for 20 min. The supernatant was collected, treated with protease inhibitor (Halt, ThermoFisher) and stored at -80 C°. Plasma total thyroxine (TT4) was quantified using a commercial ELISA kit (K050-H1, Arbor Assays, USA) with a sensitivity of 0.23 ng/mL in a standard range of 0.63-20 ng/mL. Plasma free T4 was quantified using an ELISA kit (EEL133, Invitrogen, USA) with a sensitivity of 0.94 pg/mL in a standard range of 1.56-100 pg/mL. Plasma levels of OXT were quantified using commercially available multi-species ELISA kit (K048-H1, Arbor Assays, Ann Arbor, USA) having a sensitivity of 1.7 pg/sample in a standard range of 16.38–10,000 pg/mL and an intra-assay coefficient of variation (CV) of 14%. Samples were first treated with acetone-based extraction solution followed by vacuum lyophilization. Sample concentrations were measured spectrophotometrically at 450 nm (SpectraMax 190, Molecular Devices) using standards of known concentrations. Plasma TT4, free T4 and OXT were quantified by interpolating absorbance values using a 4-parameter-logarithmic standard curve (MyAssays).

### Pup Retrieval Test

On P2-8 each litter of pups was separated from the dam for 20 min and kept on a heating pad at 35°C to maintain normal body temperature of the pups in the nest. Three randomly selected pups of either sex per litter were placed in the corners of the dam’s home cage. Maternal behaviors including latency to investigate the first pup and duration taken to return all 3 pups into the nest were scored manually using video recordings.

### Social novelty preference (SNP)

To test for the formation of short-term social recognition memory, social novelty preference (SNP) was conducted using modifications from described protocols [86]. Prior to testing days, stimulus mice were trained to stay in corrals. Mice were habituated inside an empty large cage (46 cm length × 24 width cm × 16.5 cm height) for 30 min. Mice were exposed to 2 wire interaction corrals (11 cm height × 9.5 cm inner diameter) placed at each side of the cage for 30 min. During a 5-min training trial, a sex-and age-matched stimulus mouse was placed into one corral while the empty corral was removed. After a 30 min retention period, the test mouse was introduced to the same stimulus mouse (now familiar mouse) and a novel stimulus mouse. Social recognition memory (SRM) is represented as significantly greater time spent investigating novel vs familiar stimulus as percent of total investigation time. To evaluate between-group differences, a Preference Index was calculated as the time spent investigating the trial 2 novel minus familiar stimulus / total investigation time. Using a one-sample t-test, values not greater than 0 indicated a lack of preference for novel social stimuli and abnormal social recognition ability. Test robustness was measured using a Sociability Index calculated as the ratio of time spent investigating the novel mouse in trial 1 to total investigation time during the trial 2 (**Supplementary** Fig. 2).

### Marble Burying

The marble burying (MB) test is utilized for analysis of elicited repetitive behaviors in rodents that is considered analogous to that observed in autistic individuals [87]. Exaggerated digging on MB test can provide an accurate and sensitive assay of repetitive and compulsive-like, ASD-relevant behavior in rodents since it is not correlated with other anxiety-like traits nor stimulated by novelty [88]. During MB, the test mouse was placed in the center of a polycarbonate cage (46 cm length × 24 width cm × 16.5 cm height) containing 5 cm of bedding and allowed to interact for 30 min with a 5 x 4 array of 20 equidistantly placed marbles. A minimum of ½ of the marble was defined as being buried. Mice that buried zero marbles were excluded from analysis. Bland-Altman plots indicate agreement and negligible skewing in scores provided by 2 observers (**Supplementary** Fig. 3).

### Immunofluorescence (IF) determination of OXT-ergic Neurons

Mice were sacrificed using CO_2_ inhalation and perfused transcardially with phosphate-buffered saline (PBS) followed by 4% paraformaldehyde (PFA). After 24 h post-fix, brains were cryoprotected in successive 15% and 30% sucrose solutions over 3 d. Frozen brain tissue was cryosectioned at 40 µm, mounted on gelatinized slides, and stored at −80 °C. Sections were air-dried at room temperature (RT) before being washed in PBS followed by a permeabilization/block step in 5% normal donkey serum, 1% bovine serum albumin (Vector laboratories, USA) and 0.3% Triton-X in PBS (PBS-T). Sections were then incubated with mouse monoclonal anti-OXT Neurophysin I primary antibody (PS38, 1:100; gift of Dr. Harold Gainer, NIH, USA) in PBS-T for 48 h at 4°C [89], [90]. Sections were then washed extensively with PBS followed by incubation with a donkey anti-mouse Alexa Fluor 594 secondary antibody (1:1000, Life Technologies, USA) diluted in PBS-T modified with 0.03% Triton-X at RT for 1.5 h.

Subsequent washes were followed by 1 h permeabilization/block in PBS-T containing 5% normal goat serum, 1% BSA and 0.3% Triton-X. Neuronal nuclei were then immunoprobed with rabbit anti-mouse neuron-specific nuclear protein (NeuN) (1:300, ThermoFisher) for 48 h at 4°C. Sections were then washed followed by incubation with a goat anti-rabbit Alexa Fluor 647 secondary antibody (1:1000, Life Technologies, USA) diluted PBS-T (0.03% Triton-X) containing normal goat serum for 1.5 h at RT. Sections were coverslipped with Vectashield containing DAPI (Vector Laboratories, USA).

### Microscopy and Quantitative Densitometry

For each animal, 4-8 microscope images were acquired from unilateral paraventricular (PVH) and supraoptic (SON) nuclei of the hypothalamus using a 20x/0.5objective on a Zeiss Axio Imager M2 fluorescence upright microscope (Carl Zeiss, Germany) equipped with a Hamamatsu ORCA-Flash 4.0 V3 digital camera. Images were acquired and OXT immunoreactive (ir) signal (OXT+) was quantified using densitometry software (QuPath v0.2.3, [91]). For each sample, the area of the region of interest (ROI) was traced using a mouse brain atlas and the integrated optical density (IOD) was calculated as the sum of cell area × mean intensity for the OXT+ signal and normalized to total ROI area. Results were reproduced in 3 different experiments resulting in 3–8 biological replicates per group. Representative images were taken on a Zeiss880 Airyscan Fast confocal microscope using a Plan Apochromat 20x/0.8 air objective.

### Stereological Analysis

Stereological analysis of PVH OXT and NeuN positive cells was performed as previously described, with modification [92] using a computer-assisted stereology system (Stereo Investigator, v2121.1.1, MBF Bioscience, USA). The bilateral PVH was outlined on serial sections using the DAPI channel using a live image at low magnification (10x/0.3). The Stereo Investigator placed optical disector frames in the x-y axis within each contour using systematic-random fraction sampling, outlining a 99 x 99 um sampling grid (50% of ROI sampled) with ∼15-20 counting frames for OXT and a 140 x 140 um sampling grid (25% of ROI sampled) with ∼5-8 counting frames for NeuN. The optical disector counting frames were 70 x 70 um H x W for both markers. For each mouse, a total of 3-4 sections, representing 6-8 unilateral PVHs, with a periodicity of n=1, were counted with a 40x/0.85 objective using the optical fractionator. A cell was counted when its top was in focus within the dissector height and counting frame or touching the inclusion line. Neurons touching the exclusion line were not counted. Tissue shrinkage was measured from pilot studies; the final post-processing thickness of the sections was 15 um on average. Therefore, the dissector height (Z axis) was set to 11 um to keep a top guard zone distance of 3 um. The tissue thickness was manually measured on every 4th counting frame. The optical fractionator was used to provide unbiased cell number estimates resulting in a coefficient of Error (CE, Gundersen m=1) of ≤0.1 for OXT counts and ≤0.05 for NeuN counts from all samples. All counts were made by the same investigator, who was blinded to the treatment group.

### Brain RNA extraction

Mice were sacrificed using isoflurane anesthesia and cervical dislocation. Whole brains were rapidly dissected and snap-frozen in 2-methylbutane over dry ice. Unilateral hypothalami were dissected and immediately homogenized in TRIzol Reagent (Ambion, USA) using a hand held homogenizer. Total RNA was prepared via a guanidinium thiocyanate–phenol–chloroform extraction and processed using Total RNA Miniprep Kit (New England BioLabs, USA) according to manufacturer’s instructions, modified by including DNAase1 treatment and omitting gDNA eliminator columns. Purity and quantity of RNA was assessed using nanodrop 260/280 nm and 260/230 nm ratios. Using an Agilent 2100 Bioanalyzer (Agilent Technologies Inc. USA), an RNA integrity number (RIN) >8 was deemed acceptable for downstream analysis.

### Quantitative Polymerase Chain Reaction (RT-qPCR)

RT-qPCR was used to quantitate mRNA transcripts of selected hypothalamic neuropeptides that are regulated by TH (*Oxt*, *Avp*, *Trh, Cd38*). Pre-designed DNA oligonucleotide PCR primers with efficiencies between 90 and 110% were obtained from Integrated DNA Technologies (USA) (**Supplementary Table 4**). The primer concentration ranged from 200-500nM per reaction. RT-qPCR was performed on RNA (10 ng/reaction) samples, run in triplicate, on a CFX Connect (Bio-Rad, USA) thermocycler with the Luna Universal one-step qPCR Master Mix (New England Biolabs, USA). Amplification reactions for GOIs were performed in 50 cycles of the following protocol: reverse transcription 55°C/10 min; initial denaturation 95°C/1 min; per cycle 95°C/10s denaturation, 60°C or 55°C/30s extension; 65-95°C in 0.5°C, 5s increments melt curve analysis. In each experiment, no-RNA template controls (NTCs) were run to rule out extraneous nucleic acid contamination and primer dimer formation and negative reverse transcriptase (RT). RT controls were included to rule out presence of genomic DNA. Fold-change gene expression was measured relative to the reference gene, *ActB*, and differential gene expression was determined compared to the null group using the Pfaffl method.

### Dual Immunofluorescence (IF) and Single-molecule RNA Fluorescence *in situ* Hybridization (smFISH)

At P15 or P30 offspring were sacrificed using CO_2_ inhalation and perfused transcardially with 1% diethyl pyrocarbonate (DEPC)-treated PBS followed by 4% DEPC-treated PFA. After 24 h post-fix in 4% DEPC-PFA, brains were cryoprotected in successive incubations of 10, 15 and 30% sucrose in DEPC-PBS over 3 d. Brains were embedded in OCT and stored at -80°C until further use. Brains were cryosectioned at 20 μm and sections mounted on adhesive microscope slides and stored at -80 °C. Fluorescent labeling of mRNA transcripts (RNAscope) was performed using the Multiplex Fluorescent Reagent Kit V2 Assay (ACD, USA). Sections were air-dried for 20 min before being post-fixed with 4% PFA for 60 min at 4°C, baked for 60-90 min at 60°C, dehydrated through graded ethanol solutions, treated with H_2_O_2_ and target retrieval and subjected to protease III (P30) or protease plus (P15) for 30 min at 40°C. Sections were then hybridized at 40°C for 2 h in a humidified oven with single molecule probes (ACD-Bio, USA): Mm-*Dio3*-C1 (561641), Mm-*Slc16a2*-C2 (Mct8) (545291-C2), Mm-*Thrb*-C1 (451441), Mm-*Esr2*-C2, (316121-C2), positive control (Probe-Mm-Polr2a-C1, PPIB-C2, Ubc-C3; 320881) and negative control (Probe-Mm-DapB-C1/2; 320871). Sections were incubated with Multiplex FL v.2 Amps1-3. The fluorescent signal was developed by consecutive incubation with Opal 620 for *Thrb* or *Dio3* on C1 channel (FP1495001KT), and Opal 690 for *Esr2* or *Mct8* on C2 channel (FP1497001KT, Akoya Biosciences). Sections were then immunoprobed for OXT (see above).

### Automated Quantification of smFISH and IF using Artificial Intelligence (AI) Powered Analysis Pipeline

Confocal imaging for fluorescent mRNA puncta within PVH sections was performed on an Evident FLUOVIEW FV4000 Confocal LSM at 2048x2048 16 bit resolution using a Plan-Apo 20x/0.8 air objective to generate stacks of 8-13 optical sections of 0.82-0.93 μm each. mRNA puncta were assessed using at least 4 unilateral microscope fields per animal and at least three independent animals. Channels within each image were visually inspected for background noise, artifacts, and signal integrity. Images that failed to meet quality standards were excluded from analysis. Images were then analyzed using the HALO-AI digital image analysis software (v3.6, Indica Labs, USA) to identify DAPI/OXT-stained nuclei and measure sub-cellular mRNA puncta.

HALO-AI is a trainable AI that uses deep-learning neural network algorithms to accurately segment individual cells. ROIs from each experimental group were chosen for training with HALO-AI deep learning nuclear segmentation using the Nuclei Seg V2 (DAPI) and Membrane Seg (*Oxt*) classifiers. Training was performed at a resolution of 0.32 um/px until a stable cross entropy value was observed. Classifiers were subsequently fine-tuned within the Nuclear Detection parameters to standardize identification across all treatment groups. For counts of DAPI-specific puncta, a constant 2 um was used within the cell expansion feature to allow puncta to be associated per cell. This was not necessary for OXT-positive neurons. The quality of the generated nuclear segmentation was evaluated manually by 3 experimenters. The High Plex FL (v4.1.3) module in HALO was then used for subcellular detection of mRNA transcripts.

### Statistical Analysis

Power analyses were performed to establish group size. All statistical analyses were conducted using Prism software (GraphPad 8.4.3, USA). Error bars in graphs and plots represent mean ± s.e.m, unless indicated otherwise. Sample sizes are indicated in the figure captions. Between group comparisons were accomplished using two-way or mixed model analysis of variance (ANOVA) with or without a repeated measures design or Student’s t-test to compare VEH/CON with PTU. Within groups comparisons were performed using Student’s t-test or one or two-way ANOVA if more than two groups were compared. Where the F ratio was significant, post hoc comparisons were completed using Tukey’s, Dunnet’s or Sidak’s post-hoc tests. Differences were considered statistically significant at *p*<0.05.

## Results

### Maternal intake and weight parameters

Dam food and water intake and weight gain were analyzed during gestation and lactation. We found no main effect of L-DE-71 exposure on any parameter, while H-DE-71 increased the postpartum percent absolute weight gain as compared to VEH/CON, an effect that was prevented in H-DE-71+mT4. However, the latter group showed increased relative gestational weight gain (*p*<0.01), likely due to concomitant elevation in food and water intake. PTU reduced postpartum relative food intake and postpartum absolute and relative water intake and an *apparent* augmentation of weight gain reminiscent of hypothyroidism-induced edema (**Supplementary Table 1**). PTU dams gave birth to more males (apparently high secondary sex ratio, *p*=0.055) as previously reported in rats [93], however the average number of pups per litter was not different across groups.

### Maternal behavior, plasma hormones and glycemia

Since abnormal maternal conditions may affect offspring postnatal development and adult behavior and OXT, we examined effects of DE-71 exposure on maternal pup retrieval (P2-8) and nest building behavior (P0-3). When compared to VEH/CON, DE-71-exposed dams behaved normally on a pup retrieval test that is dependent on OXTRs [94]. In contrast, PTU dams took significantly longer to locate the first pup (*p*<0.01) and also to retrieve all pups (*p*<0.05) (**Fig. 1B, C**). Moreover, mT4 caused delayed latency to locate the *first* pup, but not to retrieve *all* pups, as compared to unsupplemented corresponding groups (*p*<0.05). Mean nest building scores were only abnormal in H-DE-71 (*p*<0.05) but normalized in H-DE-71+mT4 (*p*<0.05) (**Fig. 1D**). While the PTU group mean score was within normal limits, a subgroup of dams built poor nests.

We also examined maternal plasma OXT at P22 as a proxy for central levels that are critical for maternal behavior [95]. Figure 1E shows no effects of DE-71 on mean plasma OXT levels, but a PTU-induced reduction was seen (*p*<0.05). Plasma OXT levels are dependent on TH status [96] so, as expected, mean plasma OXT was elevated with mT4 supplementation, i.e., in VEH/CON+mT4 (*p*<0.01) and L-DE-71+mT4 (*p*<0.05) but not H-DE-71, relative to unsupplemented corresponding controls, suggesting a critical effect of T4 on OXT levels.

Maternal plasma TT4 and free T4 levels were examined at P0. DE-71 produced a significant reduction in mean plasma free T4 levels at 0.1 and 0.4 mg/kg (*p*<0.05) and an apparent reduction in mean TT4 at 0.1 mg/kg (*p*=0.08, **Fig. 1F, G**). Importantly, supplementing L-DE-71 with mT4 normalized free T4 levels (*p*<0.05). As expected, PTU reduced mean plasma TT4 (*p*<0.05) and free T4 (*p*<0.001). These changes were not seen after dam manipulation had terminated at P22 (**Supplementary** Fig. 4). We also examined glycemia in exposed dams at P0-4 since TH status can affect glucose metabolism [97] and found no significant group effects (**Fig. 1H**).

### Sex-specific effects of DE-71 and mT4 on offspring brain TH content

We measured whole brain levels of pertinent TH species using mass spectrometry. Consistent with the plasma TH peak at P15 [98, 99] mean levels of TH species were 1.5-to 4-fold greater at P15 than at P30 (**Fig. 2A-P**). T4 and T3 were more abundant than rT3 and T2 as expected, since the latter are deactivation products of activity by deiodinase enzymes DIO3 and DIO2. Relative to VEH/CON, DE-71 had specific sex-and/or time-dependent effects on all TH species. In females, DE-71 exposure reduced T4 (L-DE-71, *p*<0.05) and rT3 (H-DE-71, *p*<0.05) at P15 (**Fig. 2A,C**) and elevated rT3 (L-DE-71) at P30 (**Fig. 2G**). Importantly, mT4 supplementation prevented T4 reduction, such that brain T4 in L-DE-71+mT4 was greater than L-DE-71 (*p*<0.05) and no different than VEH/CON. In P15 males, T4 was elevated by H-DE-71, T3 was elevated by L- and H-DE-71 (*p*<0.05, **Fig. 2I,J**), while rT3 and T2 were reduced by H-DE-71 relative to VEH/CON (*p*<0.05, **Fig. 2K,L**). As with females, mT4 reversed the differences observed in males (*p*<0.0001, **Fig. 2I,J**), suggesting that mT4 can prevent DE-71-evoked local TH disruption. No DE-71 exposure effects were observed in P30 males. Developmental PTU exposure reduced mean levels of all TH species in both sexes at P15 (*p*<0.01-0.0001, **Fig. 2A-D and I-L**). PTU reduced rT3 in males and females and P30 (*p*<0.05-0.0001, **Fig. 2G,O**). In P30 males PTU produced an apparent reduction in T4 (*p*=0.06, **Fig. 2M**) and a significant upregulation of T3 (*p*<0.05, **Fig. 3N**).

**Figure 2.**
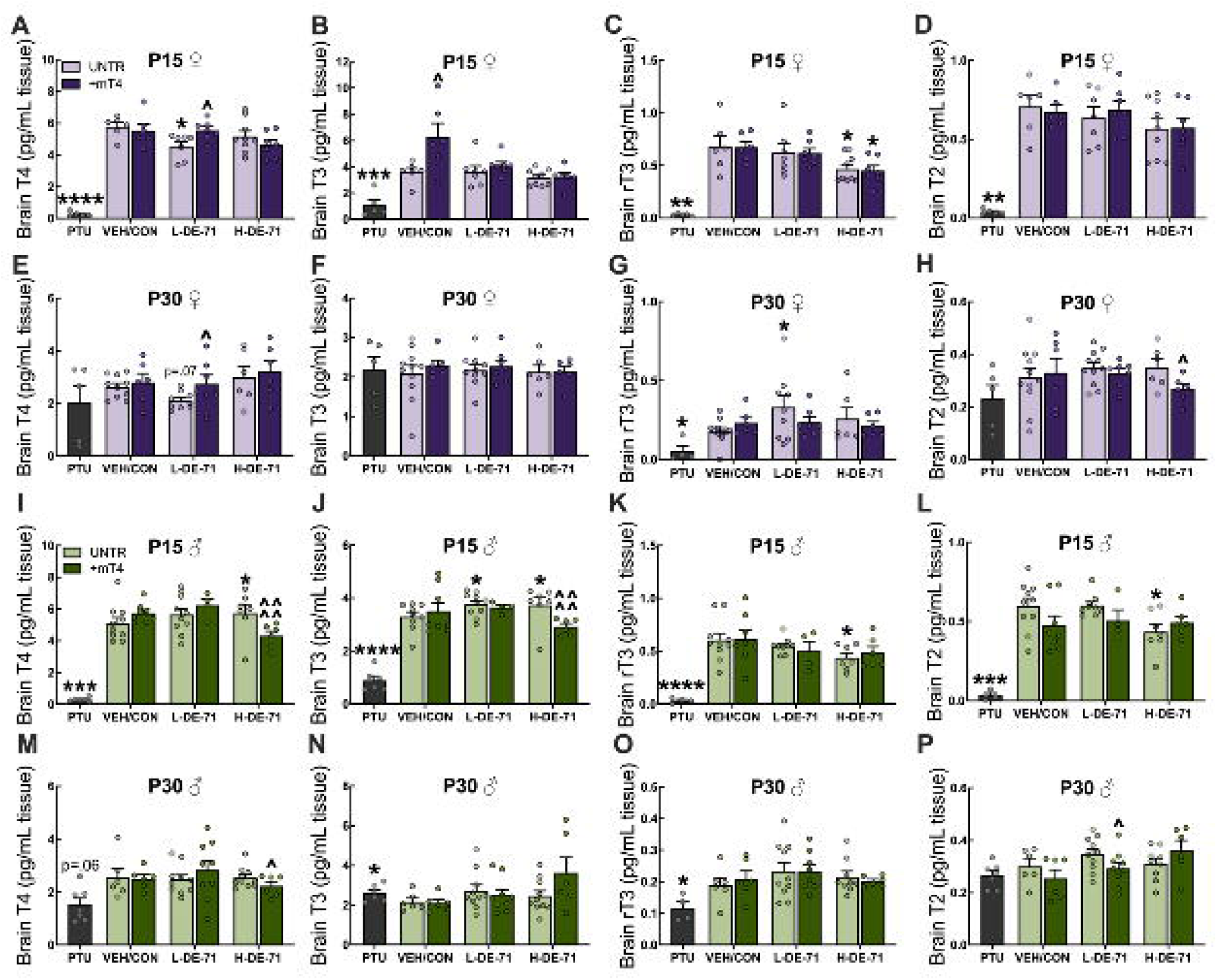
Sex-specific effects of DE-71 and mT4 on brain TH species in exposed offspring. Brain tissue concentration of TH species: (**A**) P15 female T4. (**B**) P15 female T3. (**C**) P15 female rT3. (**D**) P15 female T2. (**E**) P30 female T4. (**F**) P30 female T3. (**G**) P30 female rT3. (**H**) P30 female T2. (**I**) P15 male brain T4. (**J**) P15 male T3. (**K**) P15 male rT3. (**L**) P15 male T2. (**M**) P30 male brain T4. (**N**) P30 male T3. (**O**) P30 male rT3. (**P**) P30 male T2. Brain concentrations for rT3 and T2 in PTU-treated P15 females and males were no different than the method detection limit. *statistical difference vs VEH/CON (\**p*<0.05, \*\*\**p*<0.001, \*\*\*\**p*<0.0001). ^statistical difference vs corresponding control group (^*p*<0.05, ^^^^*p*<0.0001). *n*, 5-7/group (A-D), 5-7/group (E-H), 4-7/group (I-L), 6-10/group (M-P)

**Figure 3.**
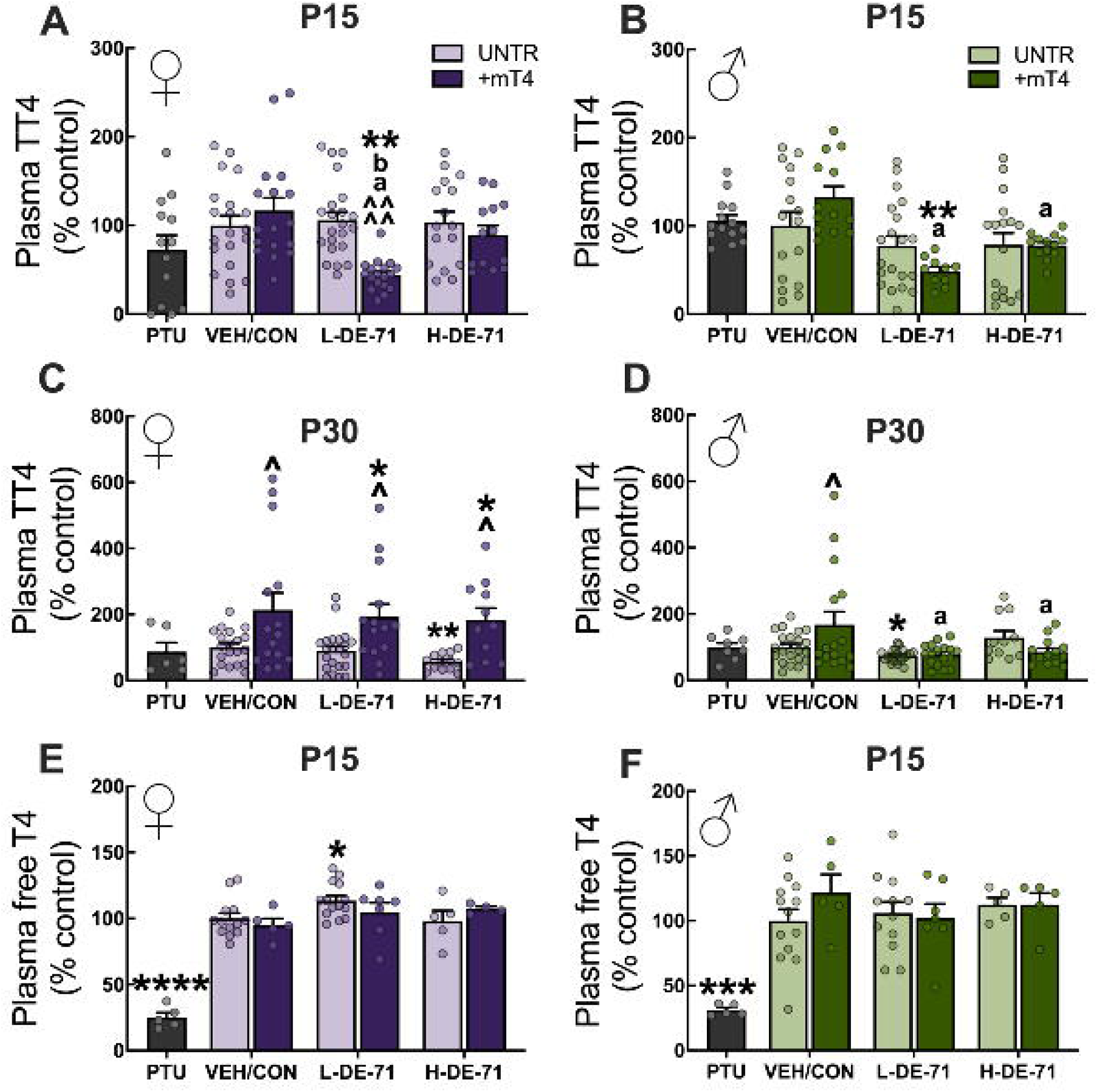
Sex-specific effects of DE-71 and mT4 on offspring plasma THs. (A,B) Plasma TT4 in P15 female (A) and male offspring (B). (C,D) Plasma TT4 concentration in P30 female (C) and male offspring (D). (E,F) Plasma free T4 concentration in P15 female offspring (E) and male offspring (F). *statistical difference vs VEH/CON (\**p*<0.05, \*\**p*<0.01, \*\*\**p*<0.001, \*\*\*\**p*<0.0001). ^statistical difference vs corresponding control group (^*p*<0.01, ^^^^*p*<0.0001). ^a^statistical difference vs VEH/CON+T4, ^a^*p*<0.05-0.0001. ^b^statistical difference vs H-DE-71+mT4, ^b^*p*<0.01. *n*, 13-22/group (A,B), 6-22/group (C,D), 4-15/group (E-F)

### Sex-specific effects of DE-71 and mT4 on offspring plasma TH

The thyroid-disrupting effects of PBDEs have been studied mostly at the plasma level in humans and rodents. DE-71 produced no effects on plasma TT4 when compared to VEH/CON at P15 (**Fig. 3A, B**). In contrast, at P30, mean plasma TT4 was reduced by H-DE-71 in females (*p*<0.01) and L-DE-71 in males (*p*<0.05), effects prevented by mT4 supplementation, indicating competition by PBDEs with THs for circulating carrier proteins. Unexpectedly, mT4 did not increase mean TT4 in P15 offspring and, instead, decreased levels when combined with L-DE-71 vs VEH/CON (*p*<0.01). Also, at P30 TT4 levels in all female mT4 groups were significantly *greater* than their corresponding controls, possibly representing a priming effect of earlier (before weaning) supplementation via the mother (*p*<0.05) (**Fig. 3C)**. Unlike P30 females, only male VEH/CON+mT4, but not male DE-71+mT4, showed elevated TT4 vs corresponding controls (*p*<0.05) (**Fig. 3D**). With regard to free T4, L-DE-71 females displayed a significant elevation (but not L-DE-71+T4) compared to VEH/CON (*p*<0.05) , suggesting displacement of bound T4, making free T4 more prone degradation and reduced brain access in L-DE-71 females but not males (**Fig. 3E, F**). Unlike DE-71 effects, PTU *reduced* mean plasma free T4 but not TT4 levels similarly in female (*p*<0.0001) and male offspring (*p*<0.001) (**Fig. 3A-F**), consistent with its antithyroid action [100]. Altogether, these results suggest time- and sex-specific effects of exposure and treatment on plasma THs.

### Maternal levothyroxine supplementation normalizes abnormal short-term social recognition memory and repetitive behavior in L-DE-71 offspring

We have previously reported a deficit in social recognition ability on a social novelty preference (SNP) test in L-DE-71-exposed female offspring [38]. Here, we show similar defects in exposed male offspring and report the rescue potential of mT4 supplementation in both sexes. Figure 4A shows significant preference for novel vs familiar conspecific in all female groups except PTU and L-DE-71 females (*p*<0.05-0.001). Importantly, L-DE-71+T4 females showed normal mean SNP scores (**Fig. 4A**). Mean Preference Index (PI) scores were not significantly different from “0” (no preference) in PTU and L-DE-71 females, using a one-sample t-test, indicating lack of social recognition ability (**Fig. 4C**). Compared to VEH/CON, PI scores were statistically lower for PTU (*p*<0.05, t-test) and L-DE-71 (*p*<0.01, 2-way ANOVA). L-DE-71+mT4 reversed PI scores compared to L-DE-71 (*p*<0.01) (**Fig. 4C**). H-DE-71 females were not deficient vs VEH/CON and showed greater SNP scores than L-DE-71 (*p*<0.05), indicating a dose-specific effect.

**Figure 4.**
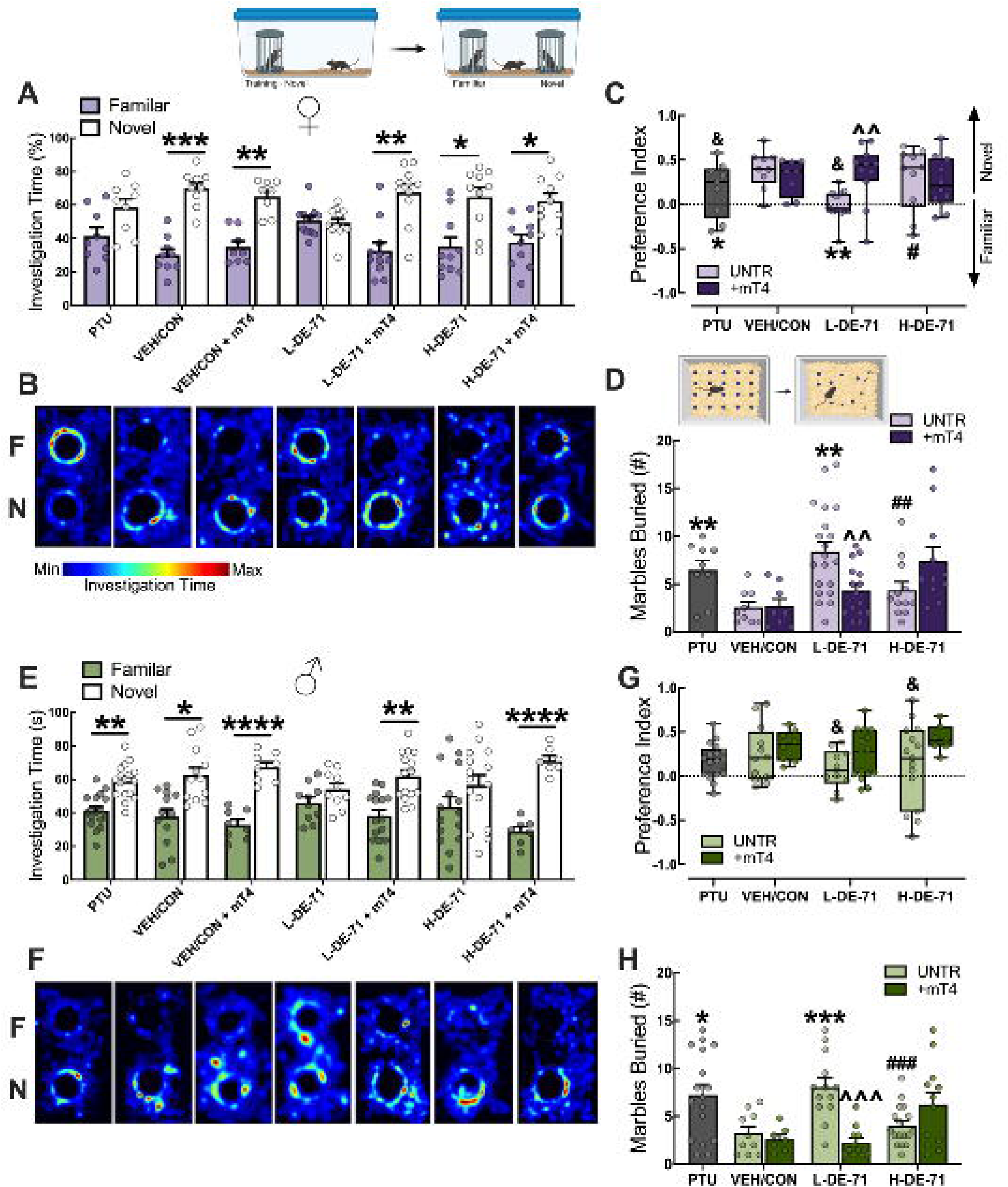
**Levothyroxine supplementation rescues deficits in social recognition ability and repetitive behavior produced by DE-71**. (**A,E**) Social novelty preference (SNP) scores in female (**A**) and male offspring (**E**). (**B,F**) Heatmaps representing investigation time in test zone for females (**B**) and males (**F**). (**C,G**) Preference Index in females (**C**) and males (**G**). (**D,H**) Number of marbles buried in a 30-minute marble burying test in females (**D**) and males (**H**). *statistical difference vs familiar stimulus in A and E (\**p*<0.05, \*\**p*<0.01, \*\*\**p*<0.001, \*\*\*\**p*<0.0001). *statistical difference vs VEH/CON in C, \**p*<0.05, \*\**p*<0.01. **^^^**statistical difference vs L-DE-71 in C,D,H, ^^*p*<0.01, ^^^*p*<0.001. ^&^lack of statistical difference vs 0 in C, G, one sample t-test; if no & symbol, *p*<0.05-0.001. ^#^statistical difference vs L-DE-71 (^#^*p*<0.05, ^##^*p*<0.01, ^###^*p*<0.001). *n*, 7-20/group

In males, DE-71, at both doses, produced abnormal social recognition ability (*p*<0.05-0.0001) (**Fig. 4E)**. As in females, mT4 normalized mean SNP scores in L-DE-71 (*p*<0.01) and H-DE-71 males (*p*<0.0001). The abnormal phenotype was verified using mean PI scores and one-sample t-tests within groups, i.e., they were not different from “0” (**Fig. 4G**). However, a two-way ANOVA did not identify statistical differences across groups, possibly indicating a weaker response to mT4 in males. Interestingly, mean SNP scores were normal in male PTU groups, in contrast to that found in females. Heatmaps for female and male offspring are shown in Figure 4B and F.

Mean scores on a marble burying (MB) test showed that L-DE-71 females (*p*<0.01) and L-DE-71 males (*p*<0.001) buried significantly more marbles relative to VEH/CON (**Fig. 4D, H**). Digging behavior was normal in H-DE-71 groups, i.e., no different than that observed in VEH/CON and significantly less than L-DE-71 in females (*p*<0.01) and males (*p*<0.001). As with SNP, mT4 prevented excessive marble burying scores in females (*p*<0.01) and males (*p*<0.001). PTU groups of both sexes displayed exaggerated burying behavior compared to VEH/CON (*p*<0.01 and *p*<0.05, respectively).

### Developmental DE-71 exposure and TH manipulation alter expression of hypothalamic neuroendocrine gene markers

Next, we examined the effects of DE-71 exposure with and without mT4 on mRNA transcripts for prosocial and related peptides in the hypothalamus. Figure 5A and B show heatmaps for DEGs with significant fold-change vs VEH/CON. In P15 females, the gene encoding thyrotropin releasing hormone, *Trh,* was down-regulated in L-DE-71 (*p*<0.05), but not L-DE-71+mT4. Also, the combination of H-DE-71 and mT4 produced downregulation of OXT (*Oxt*), *Trh,* and AVP (*Avp*) (*p*<0.05). In P30 males, *Avp* was upregulated by L-DE-71 (*p*<0.05), L-DE-71+mT4 (*p*<0.01) and by H-DE-71 (*p*<0.01). In addition, H-DE-71+mT4 upregulated *Cd38,* an enzymatic protein regulating Ca^2+^ release necessary for hypothalamic OXT release (*p*<0.01). Interestingly, PTU deregulated *Oxt* (*p*<0.05) and *Cd38* (*p*<0.01) [101]. Both OXT and CD38 are required to prevent social amnesia in social recognition tasks [68, 102, 103].

**Figure 5.**
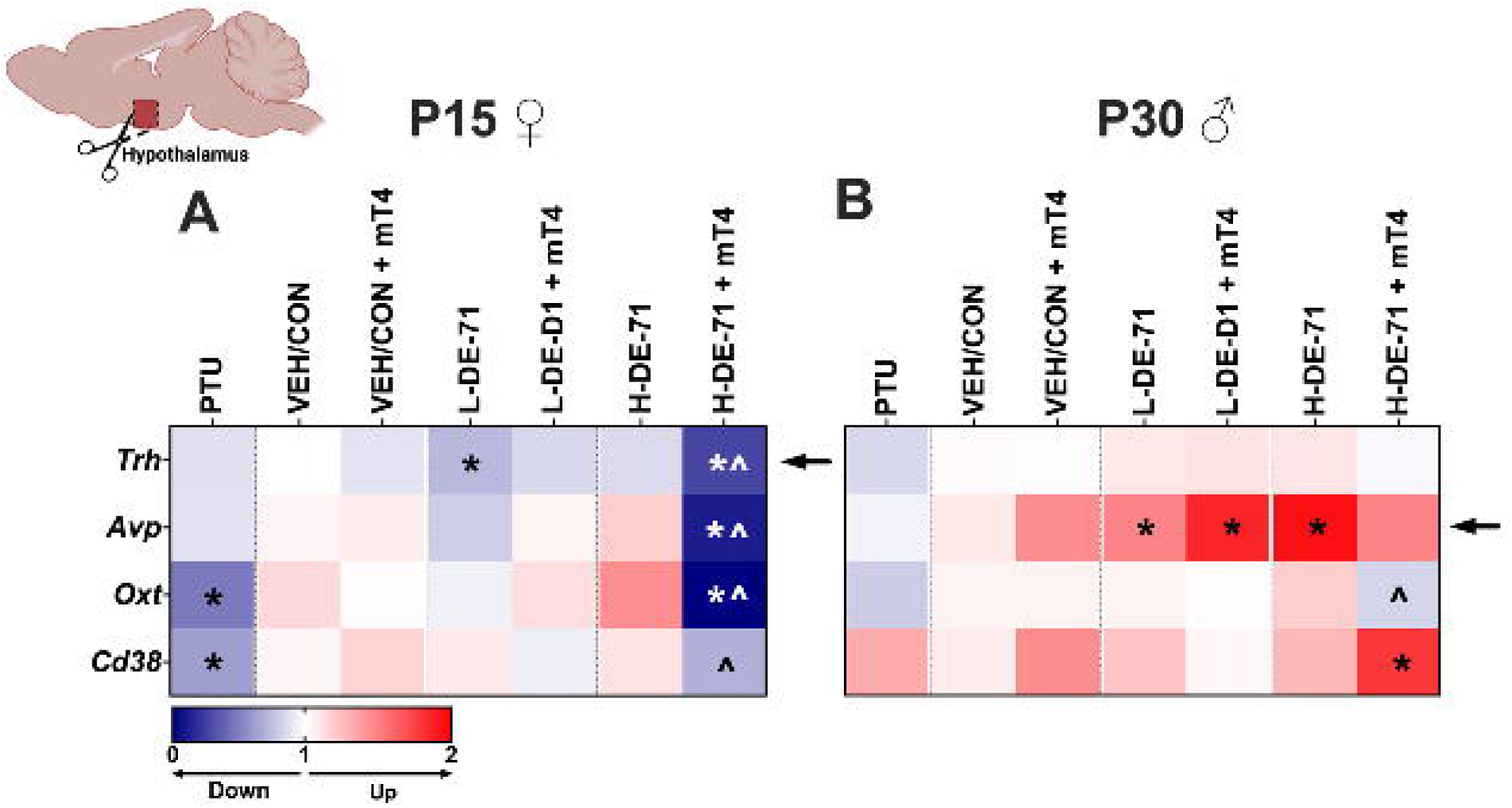
Sex-specific effects of developmental DE-71 exposure and mT4 on gene markers of prosocial neuropeptides. (A, **B)** Heatmaps of significant fold-change in relative expression of hypothalamic DEGs relative to VEH/CON in P15 females (**A**) and P30 males (**B**). Arrow indicates significant change induced by DE-71 and by additional treatment with mT4. *statistical difference vs VEH/CON, \**p*<0.05-0.0001; ^statistical difference vs corresponding control, *p*<0.05-0.0001. *n*, 3-13/group (females) and 5-10/group (males)

### Hypothalamic OXT depletion produced by DE-71 is prevented by maternal levothyroxine supplementation

Since social recognition ability is reliant on PVH OXT [104, 105], we examined changes in OXT immunoreactivity (ir) in hypothalamic sections through the magnocellular neuroendocrine nuclei, the paraventricular (PVH) and supraoptic nuclei (SON). Previous findings from our lab and others had revealed OXT as a potential target of developmental PBDE exposure in mice and rats [38, 77]. Representative micrographs in Figure 6 show large magnocellular and smaller parvocellular neurons with OXTir in the PVH. A comparison of experimental groups vs VEH/CON shows markedly reduced OXTir in DE-71 and PTU groups in females (**Fig. 6A-H**) and males (**Fig. 6L-S**). These results were corroborated using pooled data from 3-5 IF experiments. OXTir IOD was significantly reduced in L-DE-71 females (*p*<0.05) (**Fig. 6I**) and in L-DE-71 (*p*<0.05) and H-DE-71 males (*p*<0.01) (**Fig. 6T**). PTU treatment reduced OXTir IOD in both females (*p*<0.01) and males (*p*<0.01) (**Fig. 6D,I,O,T**).

**Figure 6.**
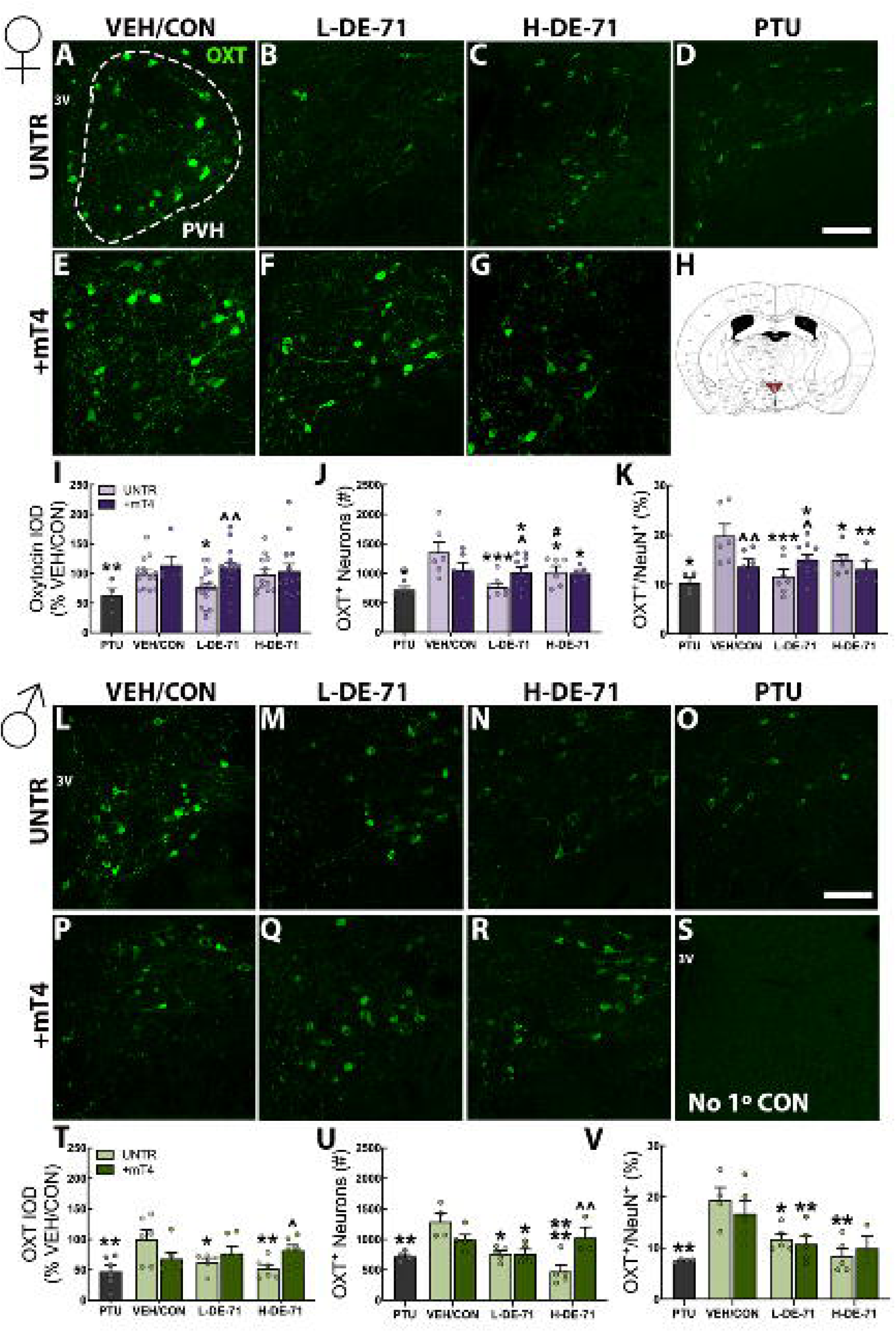
DE-71 depletes PVH OXT-synthesizing neurons in a TH-dependent manner. Representative images of OXT IF in the female (A-G) and male PVH (L-R). (H) Anatomical map of one of the 4 PVH sections used in densitometry (Bregma -.58 to -.82 mm). (S) No primary control. Quantification of OXT ir IOD in female (I) and male offspring (T). Stereological counts of OXT-neurophysin-labeled neurons in PVH in female (J) and male offspring (U). Stereological OXT^+^ neuron count as a percent of total neurons (NeuN^+^) in females (K) and male offspring (V). *statistical difference vs VEH/CON (\**p*<0.05, \*\**p*<0.01, \*\*\**p*<0.001, ****p<0.0001). ^statistical difference vs corresponding untreated control group (^*p*<0.05, ^^*p*<0.01). *n*, 4-19/group (I); 4-10/group (J,K); 5-6/group (T); 3-5/group (U,V). Scale bar, 100 um.

As a novelty of this work, stereological analysis was applied to provide a more precise measurement of PBDE effects on OXT neurons. Counts revealed fewer OXT+ neurons in both DE-71 groups of both sexes when compared to VEH/CON (**Fig. 6J,U**). To rule out potential confound due to generalized cell death, we counted the OXT+ neurons as a percent of the total neuronal population (NeuN+) and determined that specifically OXT+ neurons were reduced in L- and H-DE-71 females and males (*p*<0.05-0.001) (**Fig. 6K, V**). No group differences were seen in the number of stereologically counted NeuN+ cells, ruling out PBDE-induced cell death (**Supplementary** Fig. 5). Importantly, mT4 supplementation reversed the OXT depletion in L-DE-71 females (*p*<0.05-0.01, **Fig. 6I-K**) and prevented in H-DE-71 males (*p*<0.05-0.01, **Fig. 6T-V)**.

### DE-71 reduces male SON OXTir but not plasma levels; no rescue by maternal levothyroxine supplementation

Figure 7 shows representative micrographs of OXT-synthesizing neurons in the female (Fig. 7A**-H**) and male SON (Fig. 7K**-R**). L-DE-71 reduced OXTir IOD relative to VEH/CON in males (*p*<0.05, Fig. 7S) but mT4 treatment did not protect against this. Since the SON contains only magnocellular cells which release OXT into the systemic circulation from their axonal projections, we examined group effects on plasma OXT levels but found no effects specific to L- and H-DE-71. However, when combined with DE-71, mT4 raised plasma OXT vs VEH/CON in females (H-DE-71) and males (L-DE-71 and H-DE-71) (*p*<0.05,0.01, Fig. 7J**,T**). This may result from organohalogen interference with Ca2+ buffering mechanisms underlying neuropeptide release [75, 106]. PTU reduced mean OXTir IOD in females (*p*<0.05, Fig. 7I) and males (*p*<0.01, Fig. 7S) and elevated mean plasma OXT in females only (*p*<0.05, Fig. 7J, T).

**Figure 7.**
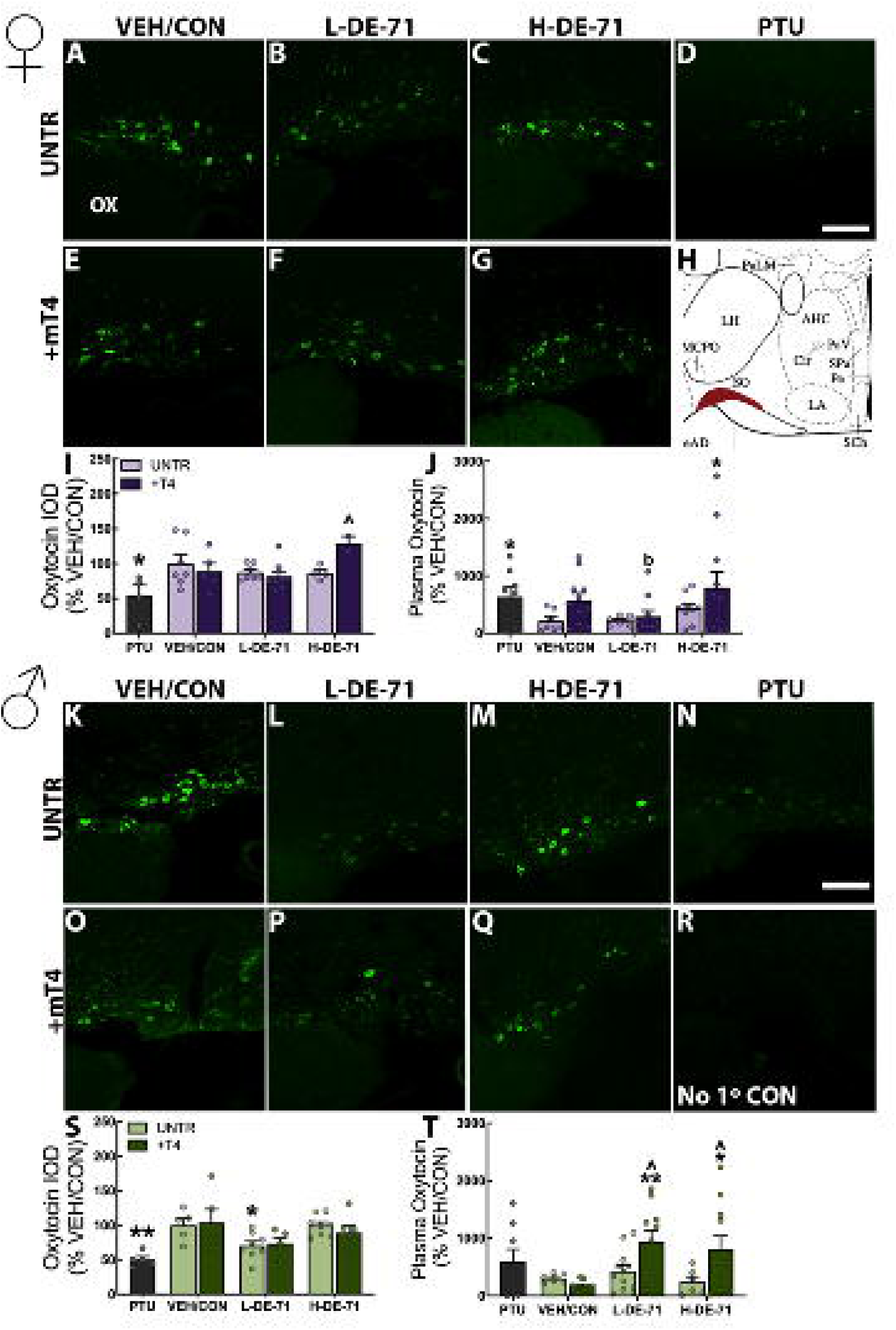
DE-71 produced depletion of SON OXT immunoreactivity (ir) in males only Representative images of OXTir in the female (A-G) and male SON (K-Q). Anatomical map of SON examined in densitometry (H; Bregma -.70 to -.82 mm). Mean values of OXTir IOD in female (I) and male offspring (S). No primary control (R). Plasma OXT in female (J) and male (T). *statistical difference vs VEH/CON (\**p*<0.05, \*\**p*<0.01). ^statistical difference vs corresponding untreated control group, ^*p*<0.05. *n*, 4-8/group (I); 7-11/group (J); 5-8/group (S); 6-12/group (T). Scale bar, 100 μm

### Sexually dimorphic deregulation of *Esr2,* but not *Thrb,* on PVH neurons

We used dual IF and multiplex RNA *in situ* hybridization to identify TH and estrogen receptors, *Thrb* and *Esr2*, that have transcriptional control of *Oxt* on OXT-synthesizing neurons, i.e., OXT*^Thrb+^*and OXT*^Esr2+^* neurons, respectively. Figure 8 shows that OXT*^Thrb+^*and OXT*^Esr2+^*neurons exhibit specific and largely non-overlapping anatomical distributions within the P30 female (**A-C**) and male PVH (**J-L**). *Thrb* was preferentially expressed in the dorsal aspect of the middle PVH which contains mostly OXT and AVP-containing magnocellular neuroendocrine cells, while *Esr2* was more visible in the center. In VEH/CON, a vast majority of OXT-ergic neurons (79, 86%) and DAPI cells (69, 75%) expressed *Thrb* in females and males, respectively (Fig. 8F**,O**). Mean PVH OXT*^Thrb+^* counts and *Thrb* counts on all DAPI-positive cells did not reveal any group differences in females (Fig. 8D**,E**) or males (Fig. 8M**,N**). For *Esr2*, nearly all VEH/CON OXT+ neurons and slightly over half of total cells showed expression in females and males (Fig. 8I, R). L-DE-71 produced selective upregulation of OXT*^Esr2+^* neurons in females (*p*<0.01, Fig. 8G) but not males (Fig. 8P). L-DE-71 deregulated *Esr2* on all DAPI-positive cells sex-dependently: upregulation in females (*p*<0.05) and *apparent* downregulation in males (*p=*0.07) (Fig. 8H**,Q**). *Esr2* expression in L-DE-71+mT4 females was similar to that found in VEH/CON, assigning mT4 a protective action from PBDE disruption. Like L-DE-71, PTU produced an *apparent* downregulation of *Esr2* selectively in male total cells (*p*=0.06; Fig. 8Q).

**Figure 8.**
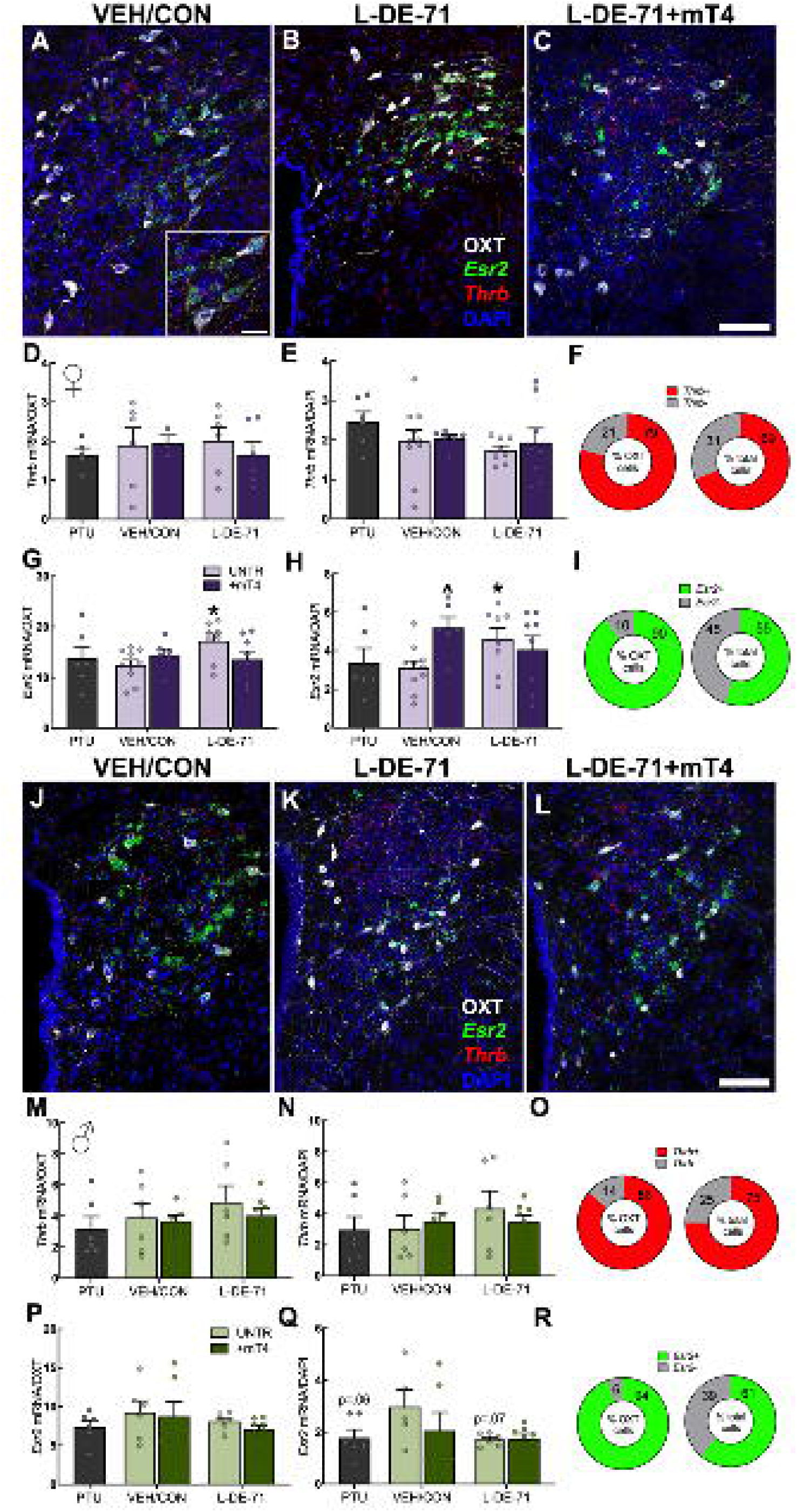
**L-DE-71 upregulates Esr2 expression in female PVH neurons**. Representative micrographs of *Thrb* (red) and *Esr2* mRNA (green) transcripts on PVH OXT- and DAPI-positive cells in P30 females (**A-C**) and males (**J-L**). Mean group values of pooled *Thrb* mRNA transcript counts in female (**D,E)** and male PVH (**M,N**). Mean group values of *Esr2* mRNA transcript counts in female (**G,H**) and male PVH (**P,Q**). PVH counts of OXT*^Mct8^* and OXT*^Dio3^* neurons as a percent of OXT+ neurons and of total cells in VEH/CON females (**F,I**) and VEH/CON males (**O,R**). *statistical difference vs VEH/CON, \**p*<0.05. **^**statistical difference vs corresponding control, \**p*<0.05. *n*, 3-10/group (D-H); 5-8/group (M-Q). Scale bar, 20 μm (A) and 200 μm (C,L)

### L-DE-71 oppositely deregulates *Mct8* and *Dio3* in female and male PVH

In comparison to *Thrb* and *Esr2* expression in PVH, *Mct8* (red) showed intermingled distribution with *Dio3* (green) (Fig. 9A**-J**). In VEH/CON a vast majority of OXT-ergic neurons (89, 99%) and DAPI cells (64, 90%) expressed *Mct8* in females and males, respectively (Fig. 9S, U). Marked group changes were observed for *Mct8* counts on OXT-producing neurons expressing *Mct8*, OXT*^Mct8+^* and all DAPI-positive cells: upregulation in P15 L-DE-71 females (*p*<0.05, Fig. 9K**,L**) and downregulation in L-DE-71 males as compared to VEH/CON (*p*<0.001-0.0001, Fig. 9O**,P**). Another TH regulatory gene, *Dio3*, was minimally expressed in female VEH/CON; few OXT-ergic neurons (10%) and DAPI cells (4%) expressed *Dio3* (Fig. 9T). Male VEH/CON had a large percentage of OXT neurons expressing *Dio3+* (71%) and most DAPI cells expressed *Dio3* (57%) (Fig. 9V). Mean counts of OXT*^Dio3+^* neurons were oppositely deregulated in female PVH (increased, *p*<0.05, Fig. 9M) and males (decreased, *p*<0.01, Fig. 9Q). These effects were not specific to OXT neurons since similar changes were observed within the PVH total cell population (**Fig 9N,R**). Importantly, mT4 supplementation prevented the L-DE-71-induced reduction in male *Mct8* (*p*<0.0001; Fig. 9O**,P**). Effects of PTU were observed as *Mct8* upregulation in female OXT neurons, OXT*^Mct8+^* (*p*<0.01), and downregulation in male DAPI-positive cells (*p*<0.05). PTU also decreased OXT*^Dio3+^* and *Dio3* on DAPI cells in males (*p*<0.01). Altogether, results from these RNAscope experiments show sexually dimorphic effects of L-DE-71 on three of the four TH-regulated mRNA transcripts studied on all PVH cell profiles. Specifically, L-DE-71 upregulated *Esr2*, *Mct8* and *Dio3* in females and downregulated *Mct8* and *Dio3* in males. mT4 ameliorated or reversed the direction of changes produced by L-DE-71 in males and females, with the exception of male PVH *Dio3* counts.

**Figure 9.**
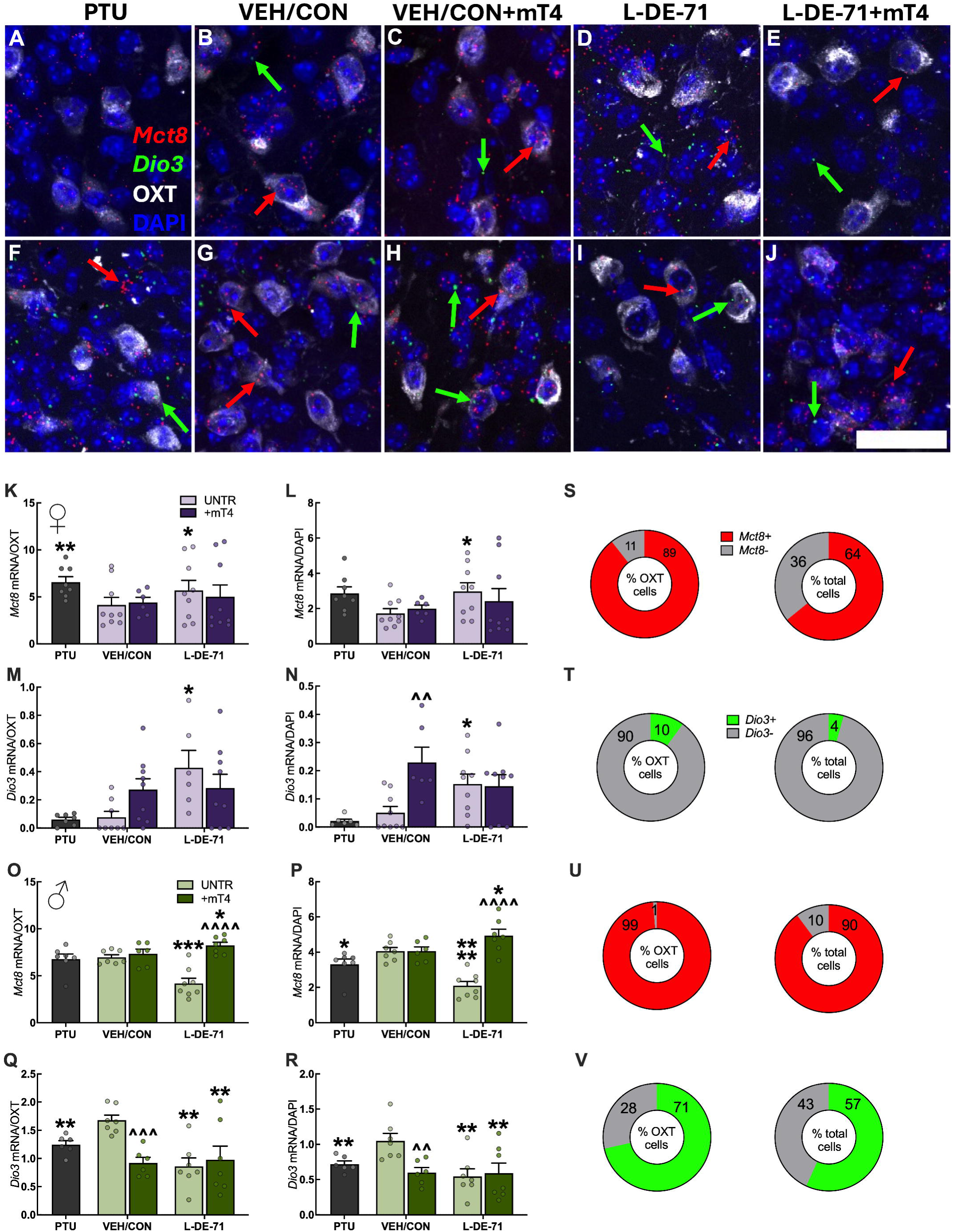
L-DE-71 oppositely deregulates Mct8 and Dio3 transcript levels in female and male PVH. **A-J:** Representative micrographs, obtained using dual IF and multiplex RNA *in situ* hybridization, of *Mct8* (red) and *Dio3* mRNA (green) on PVH OXT- and DAPI-positive cells in P15 females (**A-E**) and males (**F-J**). *Mct8* and *Dio3* mRNA distributions in PVH seemed to be co-mingled. Mean group values of *Mct8* mRNA transcript counts on OXT neurons and total DAPI+ cells in female (**K,L)** and male PVH (**O,P**). Mean *Dio3* mRNA transcript counts on OXT neurons and total DAPI+ cells in female (**M,N**) and male PVH neurons (**Q,R**). PVH counts of OXT*^Thrb^* and OXT*^Esr2^* neurons as a percent of OXT+ neurons and of total cells in VEH/CON females (**S,U**) and VEH/CON males (**T,V**). *statistical difference vs VEH/CON, \**p*<0.05, \*\**p*<0.01, \*\*\**p*<0.001, \*\*\*\**p*<0.0001. **^**statistical difference vs corresponding control, ^^*p*<0.01, ^^^*p*<0.001, ^^^^*p*<0.0001. *n*, 6-9/group (K-N); 6-8/group (O-R). Scale bar, 100 μm.

The RNAscope analysis provided OXT neuronal cell counts that served as additional information to stereological counts shown in Figure 6. A comparison of mean values for L-DE-71 vs. VEH/CON showed the same or similar reduction in PVH of females and males at P30 **(Supplementary** Fig. 6). However, at P15 L-DE-71-exposed females showed an apparent increase in the mean OXT neuron count, suggesting biphasic alterations during development in the female PVH. In contrast, OXT+ neuron counts in P15 males did not show this intermediate phenotype and instead showed a reduction.

## Discussion

The epidemic-like rise in ASD prevalence in recent decades has prompted growing concern about environmental contributors that may act alongside genetic susceptibility. Among these, PBDEs, widely used flame retardants known to disrupt endocrine function, especially in developing organisms, may act as risk factors for ASD but causative evidence is lacking. PBDEs are known to interfere with TH signaling, a pathway crucial for neurodevelopmental processes, including those that shape social behavior. Therefore, it is of critical importance to understand the endocrine-disrupting properties of flame retardants (FRs) on developing circuits and neurochemical systems underlying socioemotional behaviors fundamental to NDDs such as autism. However, to date no studies have examined the possible impact of TH-mediated disruption on the OXT system in the context of an environmental mouse model of ASD.

We reported autistic-like traits and deregulated gene markers of prosocial peptides in the SNN in adult female offspring perinatally exposed to PBDEs [38]. In the current study, we show that ASD phenotypes occur coincident with depletion of the hypothalamic OXT-ergic neuroendocrine system. We found that neurodevelopmental reprogramming caused by PBDEs is linked to sex-specific alterations in offspring brain TH status, and deregulated expression of TH-responsive genes on OXT neurons in the paraventricular nucleus of the hypothalamus, an important prosocial node in SNN [104, 105]. Importantly, we provide novel evidence that maternal levothyroxine supplementation can prevent or mitigate most PBDE outcomes, indicating that maternal TH therapy may serve to reduce risk for and/or assist in treating neurodevelopmental disorders associated with exposure to environmental toxicants. The data presented here represent the most comprehensive and detailed evaluation of PBDE exposure on the OXT system and address the novel hypothesis that PBDE exposure alters social behavioral abnormalities in the SNN via OXT in a TH-dependent manner and that reinstating OXT is sufficient to restore these ASD-relevant phenotypes.

Pregnant women and their developing offspring are particularly vulnerable to TH system disruption by PBDEs [8, 39, 107]. This is concerning since brain TH status influences brain growth and maturation during critical perinatal developmental stages. THs act on processes such as neuronal differentiation, migration, myelination and synaptogenesis and spatiotemporal precision in brain development [47, 108]. In humans, insufficient TH can induce long-term intellectual and behavioral impairments [45]. Deficiency of TH signaling results in reduced spatial learning , auditory and motor deficits and impaired maternal behavior [83, 109, 110]. Hypothyroidism, caused by maternal thyroid dysfunction, is a risk factor for childhood autism [41, 42, 44]. PBDEs, which bear structural homology to THs, can dysregulate TH status by interfering with the hypothalamic-pituitary-thyroid (HPT) axis in complex ways including TH synthesis, transport, metabolism, degradation/excretion and competition/activity with TH receptors. However, behavioral and biochemical changes have been reported in the absence of changes in circulating THs, likely resulting from altered target tissue TH status [54]. LC/MS results on brain TH species point to contrasting sex-specific levels relative to VEH/CON. L-DE-71 females showed *reduced* brain T4 at P15. This coincided with elevated plasma free T4 in females, likely due to displacement of TH from serum TH carrier proteins such as transthyretin by PBDE on mothers and offspring [39, 111]. However, this was not accompanied by changes in TT4 likely because of concurrent TH degradation/excretion that was enhanced by PBDEs [112]. PBDE-elevated brain rT3 seen at P30 may have resulted from earlier elevation in *Dio3* expression and/or reduced plasma TT4 produced by augmented hepatic TH metabolism [112, 113]. In P15 males, elevated brain T4 and T3, concomitant with reduced rT3 and T2, were dissociated from any alteration in plasma TH and may, instead, be due to local factors regulating TH status. In general, most DE-71 alterations in brain THs were seen at P15. Fewer changes were present at P30, when offspring had been weaned from mothers, and no longer received a maternal source of PBDEs. Indeed, perinatally exposed offspring show a decline in PBDE brain levels between P15 and P110 [38]. Another reason is that, after the 2nd postnatal week, endogenous TH hormonal levels have begun to normalize and their brain becomes less sensitive to TH fluctuations. Of note, P30 females born of dam mothers receiving mT4 with or without DE-71 showed a primed ability to generate plasma T4, likely representing the facilitatory effect of *in utero* mT4 on the postnatal maturation of the thyroid axis [114].

Dysregulated brain TH levels may have resulted, in part, from altered expression of genes impacting TH transport and metabolism. Evidence suggests that MCT8-mediated transport of THs and deiodinase-mediated metabolism likely dictate tissue-specific TH content [115]. Elevated brain rT3 in L-DE-71 female offspring may be linked to upregulated gene expression of DIO3, which inactivates T3 and T4 by inner-ring deiodination, that we measured in RNAscope experiments. Conversely, elevated brain T4 and T3 in DE-71-exposed males is likely the result of insufficient breakdown of T3 caused by reduced *Dio3* expression. L-DE-71 females also showed upregulated expression of TH transporter *Mct8* (also known as the *SLC16A2*), that maintains intracellular TH levels, which may represent a compensatory response to low brain T4 levels. In contrast, hypothalamic mRNA transcript levels in L-DE-71 males showed downregulated *Mct8* levels and reduced plasma TT4, a hallmark of MCT8 deficiency [116]. Importantly, mutations in the TH transporter MCT8, evident in Allan-Herndon-Dudley syndrome, a rare, X-linked genetic disorder, can produce developmental delays, intellectual disability and movement disorders [117, 118]. We also examined hypothalamic *Thrb* transcripts since nuclear T3 content and TH receptor occupancy ultimately determines the intensity of the TR-mediated signaling [117] but no group differences were observed.

Importantly, alterations in brain T4 differences produced by DE-71 in both females (L-DE-71) and males (H-DE-71), and plasma free T4 in L-DE-71 females and their dam mothers, were prevented if L-DE-71 mothers were supplemented with mT4. mT4 also prevented or mitigated alterations in hypothalamic mRNA transcripts caused by DE-71 in female (*Mct8* and *Dio3* upregulation) and male offspring (*Dio3* downregulation). Together, these results may be interpreted to mean that perinatal exposure to PBDEs dysregulated offspring brain TH status via sex-specific factors that modulate maternal and offspring TH transport and metabolism and that mT4 may work, in part, by ameliorating these pathways. Indeed, offspring rely heavily on TH transport across the placenta and blood brain barrier during prenatal and/or postnatal periods [82, 119]. We should warn that, when given to control mice with “normal” TH status, mT4 produced some adverse effects on the developing brain, such as significant reduction in the PVH OXTergic neuronal population and increased *Dio3* and *Esr2* in females and decreased *Dio3* in males (see below).

Here, we report similarly abnormal SNP and MB phenotypes, encompassing 2 major domains of diagnostic criteria for ASD, in exposed male as in female offspring [38], albeit both doses of DE-71 reduced SNP in males. Similar to PBDE outcomes reported here, developmental exposure to other FRs has been reported to disrupt various measures of social behavior in males or females (SNP, sociability, pair-bonding and sexual behaviors) [32, 33, 35, 36]. Taken together, these findings point to the influential impact of environmental toxicants, at human relevant doses, on neurodevelopment and ASD-like behavior in both sexes, especially when exposure occurs perinatally and even if exposure occurs indirectly via the mother. Furthermore, these toxicants target different brain circuits orchestrating social and repetitive behavior and may impact other ASD-relevant outcomes such as emotional recognition.

A major finding of the current study was the normalization of SNP and MB scores in DE-71-exposed offspring of both sexes if their dam mothers received levothyroxine supplementation. These results indicate that DE-71-mediated neurotoxicity is mediated via TH system disruption. Of note, mT4 protected against symptoms in 2 major domains of ASD suggesting that mT4 modulates activity of hypothalamic circuits regulating repetitive motor activity and social recognition ability [120]. In support of our findings, Tunc-Ozcan and team (2013) reported that gestational T4 treatment allowed better social recognition ability in male rat pups with fetal alcohol syndrome [121]. Another study has previously demonstrated that direct T4 treatment (15 ug/kg bw, sc) of early postnatal female rats could alleviate reference memory deficits produced by concurrent exposure to high doses of DE-71 (30 mg/d/kg bw) [122]. mT4 intervention may act, in part, by mitigating the adverse effects of PBDEs on axonal morphogenesis and oxidative stress that counteract proper neurodevelopment. This is based on a recent report showing that T3 (3-30 nM) added to BDE-47 or BDE-49-exposed primary hippocampal cultures protects axonal growth and prevents PBDE-induced ROS generation and alterations in mitochondrial metabolism [123]. Importantly, our findings support the suggested association between gestational hypothyroidism/hypothyroxinemia or congenital hypothyroidism with increased risk of autistic traits observed in epidemiological studies [41–44]. They also reveal a beneficial action of TH endocrine therapy via the adult dam mother to mitigate the autistic-like reprogramming of offspring produced by developmental exposure to environmental toxicants.

ASD-behaving offspring exposed perinatally to environmentally-relevant doses of DE-71 have depleted OXT content in the PVH and SON at P30. This finding is congruent with previous studies using the flame retardants Aroclor 1221, Firemaster 550, and another commercial PBDE mixture, DE-79, showing varied neurotoxic actions on OXT content in PVH and SON of rats and prairie voles [77, 115, 124, 125]. Our stereological analysis indicated that reduced OXT content may be due, in part, to reduced numbers of OXT-ergic neurons in the PVH; the development and survival of this neuronal population is crucial for social recognition and social environment selection [126]. Through their neuronal projections to specific forebrain nuclei, PVH OXT+ neurons can regulate sociosexual behavior [66, 104, 105, 127]. We speculate that DE-71-induced depletion of OXT compromises the capacity for PVH-mediated OXT signaling leading to deficient social recognition ability. It is likely that the reduced OXT signaling is similarly disrupted as the diminished vasopressin (AVP) release observed in magnocellular neuroendocrine cells following in vitro DE-71 exposure, potentially due to interference with calcium-dependent exocytosis and neurotransmitter release mechanisms [76]. Indeed, DE-71 effects on PVH OXT depletion in females and males were characterized with the same dose dependence as that producing their aberrant social recognition: low dose (0.1 mg/kg) in females and low and high dose (0.4 mg/kg) in males. In support of our interpretation, social cues activate the PVH OXTergic population and reducing the number of PVH-OXT neurons produces abnormal sociability behavior in an OXTR-dependent manner [104]. Specific silencing of PVH OXT+ neurons leads to an impairment in short- and long-term social recognition memory [105]. Earlier studies have supported the importance of OXT signaling in social recognition memory by demonstrating that OXT or OXTR knockout mice investigate conspecifics with no preference for novel over familiar mice [68, 128]. Reduced hypothalamic OXT may also explain the excessive repetitive behavior observed in our L-DE-71 mice since acute administration of OXT decreases repetitive stereotypical behaviors in males in ASD studies [129]. Maternal attachment and gut microbiota influence offspring PVH OXT neuronal counts and OXT peripheral levels as well as social novelty preference since they are all compromised if mouse litters are cross-fostering with non-biological mothers [130].

Another novelty of our study is that maternal levothyroxine supplementation prevented reduction in PVH OXT content, observed in L-DE-71 female and H-DE-71 male group. This recovery effect of mT4 is remarkable for three reasons. First, it suggests that PBDEs actions in promoting ASD-like behavior and deleting hypothalamic OXT are linked. Second, it suggests that the ASD-like traits produced by developmental PBDEs are possibly due to neuroendocrine disruption of prosocial OXT-mediated signaling via TH system disruption. This is consistent with the regulatory action of T4 on the OXT and OXTR mRNA promoter [70] and on plasma OXT in female mice and male rats [96, 131]. Furthermore, maternal TH status might be developed into a novel and much-needed biomarker for some cases of ASD *in utero.* Indeed, there is a strong association between severe maternal hypothyroidism and risk of ASD [41, 42, 44, 132]. Given the preventive effects of maternal levothyroxine treatment against toxic actions of PBDEs and possibly other EDCs, further study is warranted to determine the effectiveness and safety (timing and dosage) in humans.

TH action depends on regulatory regions of responsive genes known as TH responsive element (TRE) and estrogen responsive element (ERE) that are binding sites for TH receptors (TR) and estrogen receptors (ER) [133]. Both TR and ER can activate TRE and regulate TH responsive genes independently, but ER inhibits TRE activity when TR is present. Using dual OXT IF and multiplex RNA *in situ* hybridization we found no PBDE effects on TR (*Thrb*). In contrast, L-DE-71 upregulated mean mRNA transcripts (*Esr2*) of estrogen receptor beta (ERß) in OXT*^Esr2+^* neurons of females. OXT production in the PVH and OXTR receptor density in various extrahypothalamic areas that are part of the SNN are regulated by endogenous estrogen acting on ERB [134, 135]. From knockout, pharmacological and optogenetic studies we have learned that these substrates are critical for social recognition ability, making ER relevant to the deficiencies in PVH OXT content and social behavior shown here [66, 135–137]. Others have reported female-specific *Esr2* deregulation in PVH by gestational PCB exposure [124]. Together with our current findings, *Esr2* can be identified as a common target of flame retardant EDCs. PBDE congeners in DE-71 may regulate ERB gene expression via their activity as antagonists and agonists of ERB [37]. L-DE-71 also upregulated *Esr2* transcripts in all female PVH cells, the majority of which express ERB in the rodent PVH [138]. This hypothalamic target of PBDEs may affect other regulatory functions associated with the PVH, some of which occur in a sexually dimorphic manner.

Our molecular examination of *Dio3*, the TH-regulatory gene *Dio3,* the gene encoding iodothyronine deiodinase 3, which regulates TH levels and limits their biological action [139], revealed that L-DE-71 differentially deregulated OXT*^Dio3+^* neurons in exposed females (up) and males (down). In support of our findings, upregulated *Dio3* (in hippocampus) has been correlated with compromised social memory due to gestational ethanol exposure [121]. However, it is unclear if downregulated *Dio3* may also be influential in deficient social recognition in L-DE-71 males. For example, *Dio3* deletion does not impact sociability in females nor males, although it produces aggression-related behavior and abnormal neonatal/adult expression of *Oxt, Avp* and their receptors [131]. L-DE-71 also produced sex-specific changes in hypothalamic expression of *Mct8* (see above). The combined results from these RNAscope experiments show sexually dimorphic effects of L-DE-71 on three of the 4 TH-regulated mRNA transcripts studied. Specifically, L-DE-71 impacted OXT^Esr2+^ and OXT*^Dio3+^*neurons in females, both of which were upregulated, and OXT*^Mct8+^*and OXT*^Dio3+^* neurons in males, both of which were downregulated. Notably, mT4 prevented or mitigated all deregulated molecular changes except *Dio3* downregulation in males. Based on our findings, we believe that the autistic traits displayed in PBDE-exposed females and males may result from targeting of distinct TH-responsive molecular elements on OXT and other PVH neurons equally leading to abnormal TH status, OXT depletion.

The processes of biosynthesis and release of OXT (and AVP) can be modulated by thyrotropin-releasing hormone (TRH) [140]. Our PCR data showed a female-specific reduction of hypothalamic *Trh* expression caused by L-DE-71 at P15, suggesting an altered regulation of OXT by TRH. Transcriptional regulation of *Trh* by PBDEs may result indirectly by increased hormonal feedback resulting from elevated plasma free T4 in females and/or by direct action on *Trh*-expressing PVH neurons. Transcriptional alterations in the male hypothalamus were also evident as significant mRNA elevation with both doses of DE-71. Together, these results reveal sex-specific changes in TH-responsive regulators of prosocial neurochemical systems targeted by DE-71 that may indicate different underlying mechanisms for aberrant social behavior. Taken together, the changes to protein and gene expression caused by maternal transfer of PBDEs to offspring during their development suggest that transcriptional and posttranscriptional mechanisms play roles in mediating how PBDEs reprogram hypothalamic development.

The neurotoxic actions of DE-71 were not identical to those of the anti-thyroid drug PTU suggesting divergent TH-disruptive actions, especially in developing offspring. In contrast to the more specific actions by DE-71, PTU reduced all brain TH species in offspring of both sexes, most markedly seen at P15, consistent with blockade of TH production and its extrathyroid inhibition of deiodinase conversion of T4 to T3 [141]. Maternal PTU on P30 offspring plasma and brain had mostly but not entirely disappeared. Lingering adverse effects of PTU on brain neurogenesis have been reported up to 6 weeks post exposure [142]. Both doses of DE-71 acted similar to PTU in disrupting dam plasma THs (reduced plasma free T4 and TT4) as reported previously [54] while, in offspring, DE-71 and PTU produced differential effects, decreased TT4 at P30 or decreased plasma free T4 in P15 in female and male offspring, respectively. Developmental PTU treatment reduced SNP scores in a manner similar to L-DE-71, but only in female mice, while PTU affected male and females similarly on the MB test. PBDE-induced behavioral deficiencies in offspring are not likely due to alterations of maternal physiology or behavior. In contrast, PTU reduced dam scores on maternal attachment test, an effect that may be attributed, in part, to reduced plasma OXT observed in dams [143]. PTU also reduced gestational food and water intake and apparently augmented postpartum weight gain, changes reminiscent of profound maternal hypothyroidism.

## Conclusions

For the first time we have shown that maternal transfer of the environmentally relevant mixture of PBDEs resulted in a reduction of PVH OXT-ergic neurons coincident with deficient social recognition memory and exaggerated repetitive behavior at P30. While these ASD-relevant phenotypes were seen in exposed offspring of both sexes, females were mostly affected by 0.1 mg/kg while male offspring were equally susceptible to 0.1 and 0.4 mg/kg doses of DE-71. Another novelty of our findings is that PBDE outcomes occurred in a TH-dependent manner since levothyroxine supplementation in exposed dam mothers prevented or significantly mitigated the abnormalities in offspring behavior, hypothalamic OXT neurochemistry and expression of genes encoding TH metabolism (*Dio3*), and transport (*Mct8*) and co-regulation factors (*Esr2*) on PVH cells. In support of the thyroid-disrupting properties of DE-71, THs regulate pro-social hormones such as OXT and AVP, which are critical for social recognition ability and other ASD co-morbidities. In our hands, DE-71 altered TH status in an age- and sex-specific manner, by affecting circulating and/or brain THs. Taken together, this study provides affirmative evidence for the novel hypothesis that environmental toxicants contribute to autism phenotypic behavior by depleting central OXT signaling via TH system disruption and that maternal levothyroxine treatment may prevent expression of these neurodevelopmental aberrations. Further, our findings point to TH-dependent transcriptional mechanisms occurring in a sexually dimorphic manner. Based on these and existing human data, novel therapies focused on diagnosis and management of maternal TH status may be beneficial in mitigating environmentally altered regulation of offspring TH and OXT signaling that is implicated in childhood autism, social amnesia and schizophrenia [65, 68, 144].

### Limitations of Study and Cautionary Statement

One important interpretation of our findings is that DE-71 induces ASD-relevant behavioral phenotypes through its OXT-depleting effects, but whether social deficits in ASD cause low OXT levels or *vice versa* is unclear. Our PVH OXT cell counts at P15 and P30 are consistent with disruption during critical windows of OXTergic development and help support a causative role for early OXT system disruption in PBDE outcomes. These alterations may be linked to PBDE-compromised local TH status as supported by changes in TH regulatory neuromolecular markers in the hypothalamus. Nevertheless, associated data on TH status such as whole brain TH LC/MS and plasma T4 immunochemical measurements represent broader indices that may be insufficient, especially since T3 or TSH measures were lacking. Here, we interpret PBDE-induced OXT depletion in the context of social behavior but PBDEs may also impact other PVH OXT-regulated functions such as lactation, stress, hydromineral balance and autonomic regulation.

Finally, while our findings are novel and may inspire compelling hypotheses, the observed behavioral and gene expression patterns are correlational, and their causal relationship remains to be explored in future research.

Importantly, maternal T4 (mT4) supplementation prevented both OXT neuron depletion and ASD-like behaviors in offspring, highlighting its therapeutic potential. This is relevant given that children with ASD have been reported to exhibit lower serum OXT [145], and confirmation from animal studies that deficient OXT or AVP signaling can mimic autistic traits [146, 147]. While intranasal OXT has shown limited clinical efficacy, our data indicate that upstream targeting of the central OXT system via TH manipulation may add benefit. Gestational hypothyroidism, especially severe TH deficiency, is a known ASD risk factor [40, 44], but TH supplementation is not currently used for autism treatment. Controversies surrounding treatment of mild maternal hypothyroidism are based on evidence of preterm birth [148], but treatment can prevent preterm birth, gestational hypertension, placental abruption, and neonatal death in more severe hypothyroid cases. Existing clinical trials with T4 therapy show promise, particularly when given to extremely premature infants to improve motor, language and cognitive scores at four years [149], though more data is needed to justify a change in clinical practice. Clinicians warn that maternal iodine and TH levels should be managed to ameliorate the risk of ASD [42]. Our findings suggest that suppressing toxicant burden, in part via mT4, may provide benefit too. Levothyroxine, an FDA-approved therapy for underactive thyroid diseases in children [117], could be repurposed for early intervention in at-risk pregnancies, though caution is warranted due to potential adverse effects with overdosing. Indeed, we found that mT4, when given to *unexposed* vehicle controls, altered PVH OXT neurons and gene expression, suggesting that dosing and developmental timing must be individually optimized.

## Acknowledgements

We thank Dr. Harold Gainer (deceased) for gift of Neurophysin antibodies. We thank Drs. R. Hartman (Loma Linda University) for use of Ethovision. We thank Alan Gilman from Evident Scientific for assistance with imaging. We are grateful to Drs. Levi Maston and Scott Rowlins (Indica Labs) for help with HALO software and to Dr. A. Gupta (UC Riverside) for his helpful comments on T4 supplementation. We are grateful to Drs. N. zur Nieden, F. Sladek and I. Ethell (UC Riverside) lab for the gift of mice and to Y. Korde for technical assistance. Images were created with Biorender.com.

## Declarations Disclaimer

Research reported in this publication was supported by NIEHS (F31ES034304). The content is solely the responsibility of the authors and does not necessarily represent the official views of the National Institutes of Health.

## Ethics approval

Care and treatment of animals was performed in accordance with guidelines from and approved by the University of California, Riverside Institutional Animal Care and Use Committee (AUP# 20170026, 20200018 and 5).

## Consent to participate

Not applicable.

## Consent for publication

All authors reviewed and approved the final manuscript.

## Availability of Data and Material

Not applicable. **Code Availability** Not applicable.

## Permission to reproduce material from other sources

Not applicable.

## Author Contributions

**Conceptualization,** E.V.K., K.-W.S., M.C.C.-C.; **Methodology**, E.V.K., M.E.D., M.D.A., K.-W.S., M.C.C.-C.; **Validation**, E.V.K., M.E.D., K.-W.S., M.C.C.-C.; **Formal Analysis**, E.V.K., M.E.D., A.E.B., M.D.A., M.C.C.-C.; **Investigation**, E.V.K., M.E.D., A.E.B., C.N.L., M.D.A., L.C., A.H., A.A.L., A.G.C., N.L., D.C., T.M., K.-W.S., M.C.C.-C.; **Writing – Original Draft**, E.V.K., M.C.C.-C.; **Writing – Reviewing and Editing**, E.V.K., M.E.D., A.E.B., C.N.L., M.D.A., A.A.L., M.C.C.-C.; **Visualization**, E.V.K., M.E.D., A.E.B.; **Resources**, E.V.K., T.M., K.-W.S., M.C.C.-C.; **Data Curation**, E.V.K., M.E.D., A.E.B., M.D.A.; **Supervision**, E.V.K., M.C.C.-C.; **Project Administration**, E.V.K., M.C.C.-C.; **Funding Acquisition**, E.V.K., T.M., K.-W.S., M.C.C.-C. All authors reviewed and approved the final manuscript.

## Abbreviations

ACTB: beta-actin
AI: artificial intelligence
ANOVA: analysis of variance
ASD: Autism Spectrum Disorder
AVP: vasopressin
BFR: brominated flame retardants
bw: body weight
CON: control
DAPI: 4′,6-diamidino-2-phenylindole (nuclear stain)
DE-71: commercial PBDE mixture (penta brominate diphenyl ether)
DIO: deiodinase
DMSO: dimethyl sulfoxide
EDC: endocrine-disrupting chemical
ELISA: enzyme-linked immunosorbent assay
ERE: estrogen response element
ERβ: estrogen receptor beta
ESI: electrospray ionization
FRs: Flame Retardants
GD: gestational Day
G x E: gene-by-environment interactions
GOI: gene of interest
HALO-AI: deep learning digital image analysis platformed by Indica Labs
H-DE-71: high dose DE-71 group (0.4 mg/kg bw/day)
HPT: hypothalamic-pituitary-thyroid
IF: immunofluorescence
IOD: integrated optical density
ir: immunoreactive
IACUC: institutional animal care and use committee
LC/MS: Liquid chromatography–mass spectrometry
L-DE-71: low dose DE-71 group (0.1 mg/kg bw/day)
LSM: laser scanning microscope
MCT8: monocarboxylate transporter 8
MDL: method detection limit
MNC: magnocellular neuroendocrine cell
mRNA: messenger ribonucleic acid
mT4: maternal levothyroxine supplementation
MRM: multiple reaction monitoring
NDD: neurodevelopmental disorder
NeuN: neuronal nuclei marker
NTC: no-RNA template control
OXT: oxytocin
PBS: phosphate-buffered saline
PBS-T: PBS with Triton-X
PBDEs: polybrominated diphenyl ethers
PFA: paraformaldehyde
PI: preference index
PND: postnatal day
POP: persistent organic pollutants
PTU: propylthiouracil
PVH: paraventricular nucleus of the hypothalamus rT2 - 3,5-diiodo-L-thyronine
rT3: reverse 3,3’,5’-triiodothyronine\
RIN: RNA integrity number
RNA: ribonucleic acid
ROI: region of interest
RT-qPCR: reverse transcription quantitative polymerase chain reaction
SNN: social neural network
Sm FISH: single-molecule fluorescence *in situ* hybridization
SNP: social novelty preference
SRM: social recognition memory
SRS: social responsiveness scale T1 - 3-iodo-L-thyronine
T1AM: 3-iodothyronamine T2 - 3,3′-diiodo-L-thyronine T3 - 3,3′,5-triiodothyronine T4 - L-Thyroxine
TH: thyroid hormone
TRE: thyroid hormone response element
TRH: thyrotropin-releasing hormone
TSH: thyroid stimulating hormone
TT4: total T4 UNTR: untreated (without mT4) VEH/CON - vehicle control group

