## Supplementary Information for "Maternal Thyroid Supplementation Prevents Autistic-relevant Social Behavior and Hypothalamic Oxytocin Depletion Produced by Developmental Exposure to Environmental Toxicants"

### Supplementary Table 1

### Supplementary Table 1. Dam Physiological Parameters during Gestation and Lactation

| | PTU | VEH/CON | VEH/CON+T4 | L-DE-71 | L-DE-71+T4 | H-DE-71 | H-DE-71+T4 |
|---|---|---|---|---|---|---|---|
| Maternal Parameters | | | | | | | |
| n[i] | 7-12 | 7-17 | 4-12 | 7-12 | 6-15 | 5-13 | 7-14 |
| **Gestational food intake** (GD15-0) | | | | | | | |
| Absolute (g/d) | 7.67±0.74 | 7.00±0.67 | 8.88±0.70 | 7.42±0.37 | 6.98±0.33 | 6.28±0.35 | 8.87±1.45 |
| Relative (g/d/pup) | 0.96±0.12 | 0.94±0.09 | 1.45±0.18 | 0.99±0.07 | 1.16±0.11 | 0.97±0.15 | 1.76±0.37*^ |
| **Postpartum food intake** (P1-P10) | | | | | | | |
| Absolute (g/d) | 12.90±1.07 | 15.59±1.02 | 12.13±0.95 | 14.94±1.15 | 13.02±0.71 | 11.59±0.43 | 13.36±1.24 |
| Relative (g/d/pup) | 1.61±0.10* | 2.10±0.13 | 2.09±0.18 | 1.98±0.14 | 2.09±0.14 | 1.82±0.18 | 2.28±0.22 |
| **Gestational water intake** (GD15-P0) | | | | | | | |
| Absolute (mL/d) | 8.10±0.52 | 7.00±0.35 | 7.63±0.46 | 7.76±0.19 | 7.02±0.18 | 7.31±0.30 | 7.56±0.30 |
| Relative (mL/d/pup) | 0.99±0.08 | 0.96±0.07 | 1.32±0.16 | 1.05±0.06 | 1.17±0.13 | 1.08±0.11 | 1.53±0.20* |
| **Postpartum water intake** (P1-P10) | | | | | | | |
| Absolute (mL/d) | 12.79±0.44* | 14.46±0.48 | 13.62±0.66 | 14.12±0.45 | 13.90±0.33 | 13.47±0.40 | 12.47±0.78 |
| Relative (mL/d/pup) | 1.58±0.09* | 1.91±0.12 | 2.28±0.15 | 1.88±0.11 | 2.24±0.25 | 2.04±0.20 | 2.30±0.28 |
| **Gestational weight gain** (GD15-P0) | | | | | | | |
| Absolute (g/d) | 1.85±0.21 | 1.90±0.11 | 1.58±0.20 | 1.58±0.14 | 1.59±0.17 | 1.64±0.19 | 2.30±0.21 |
| Relative (g/d/pup) | 0.23±0.02 | 0.22±0.03 | 0.26±0.03 | 0.22±0.02 | 0.24±0.03**b** | 0.27±0.04 | 0.41±0.05** |
| Abs. Percent Change (%)[ii] | 41.19±6.09 | 52.80±4.28 | 44.38±6.50 | 50.34±4.72 | 48.28±5.28 | 50.56±6.00 | 52.91±3.51 |
| Rel Percent Change (%) | 4.94±0.46 | 6.74±0.67 | 7.41±1.04 | 6.98±0.75 | 7.50±0.93 | 8.57±1.37 | 8.94±0.73 |
| **Postpartum weight gain** (P1-P10) | | | | | | | |
| Absolute (g/d) | 0.61±0.20 | 0.22±0.11 | 0.11±0.20 | 0.58±0.06 | 0.27±0.11 | 0.78±0.21**a** | 0.58±0.13 |
| Relative (g/d/pup) | 0.08±0.03 | 0.03±0.02 | 0.01±0.03 | 0.08±0.01 | 0.04±0.02 | 0.09±0.02 | 0.08±0.02 |
| Abs. Percent Change (%)[iii] | 10.66±3.79 (*p*=0.12) | 3.51±2.07 | 2.51±5.25 | 10.95±1.27 | 5.25±2.06 | 17.22±4.24***a** | 11.21±2.12 |
| Rel. Percent Change (%) | 1.33±0.42 (*p*=0.13) | 0.50±0.28 | 0.27±0.92 | 1.47±0.25 | 0.73±0.30 | 1.95±0.43 | 1.56±0.28 |

| | | | | | | | |
|---|---|---|---|---|---|---|---|
| **Number of Pups in Litter** | 7.50±0.58 | 8.00±0.44 | 6.82±0.62 | 7.64±0.45 | 7.00±0.49 | 7.23±0.70 | 6.43±0.68 |
| **Secondary Sex Ratio** (males/males+females) | 0.63±0.04 (*p*=.055) | 0.49±0.05 | 0.49±0.07 | 0.44±0.04 | 0.51±0.06 | 0.48±0.07 | 0.55±0.05 |

Maternal food intake was measured from GD15-P10. Relative food intake was normalized to total pups in litter (relative). Gestational weight gain was calculated as difference in weight from pre-pregnancy to late gestation (GD 15-P0). Relative weight gain was normalized to pre-pregnancy weight (initial) and to total pups in litter (relative). Gestational weight gain relative to initial weight was calculated as percent change from weight at GD 2-3 (whichever was 18d prior to birth) until birth (usually GD19). Data are expressed as mean±s.e.m.

[i]Indicates number of dams/treatment group

[ii]Change vs initial weight at 18d before pup birth

[iii]Mean change per day vs initial weight at P1

*Statistical difference vs VEH/CON, *$p<0.05$

^Statistical difference with corresponding mT4 group, ^$p<0.05$

[a]Statistical difference vs VEH/CON+mT4, [a]$p<0.05$

[b]Statistical difference vs H-DE-71+mT4, [b]$p<0.05$

**Supplementary Statistical Information for Supplementary Table 1**

Gestational food intake was measured from GD15 until birth (P0). Gestational absolute food intake differed based on T4 treatment (Two-Way ANOVA: *Treatment Effect* $F_{(1,47)}=5.29$, $p=.025$), but there were no differences on post-hoc comparisons. No differences were observed across exposure or interaction (*Exposure Effect* $F_{(2,47)}=0.54$, ns; *Treatment x Exposure Interaction* $F_{(2,47)}=2.67$, ns). No difference was seen between PTU and VEH/CON groups (Independent *t*-test: t=0.67, *ns*). Gestational relative food intake, defined as absolute food intake per number of pups per dam, differed based on T4 treatment (*Treatment Effect* $F_{(1,46)}=11.45$, $p=.002$), with H-DE-71+T4 dams showing significantly greater relative food intake compared to VEH/CON ($p=.018$), and H-DE-71 ($p=.037$). No differences were observed across exposure or interaction (Two-Way ANOVA: *Exposure Effect* $F_{(2,46)}=1.45$, *ns*; *Treatment x Exposure Interaction* $F_{(2,46)}=1.69$, ns). No difference was seen between PTU and VEH/CON groups (Independent *t*-test: t=0.10, *ns*).

There were no group differences in postpartum absolute food intake from P1-10 (Two-Way ANOVA: *Treatment Effect* $F_{(1,50)}=2.01$, *ns*; *Exposure Effect* $F_{(2,50)}=1.24$, ns; *Treatment x Exposure Interaction* $F_{(2,50)}=3.15$, *ns*). No difference was seen between PTU and VEH/CON groups (Independent *t*-test: t=1.75, *ns*). Likewise, postpartum relative food intake was not different across groups (Two-Way ANOVA: *Treatment Effect* $F_{(1,50)}=1.94$, ns; *Exposure Effect* $F_{(2,50)}=0.01$, *ns*; *Treatment x Exposure* $F_{(2,50)}=1.01$, *ns*). Relative food intake was higher in VEH/CON compared to PTU (Independent *t*-test: t=2.69, $p=.016$) (**Supplementary Table 1**).

Gestational water intake was also measured from GD15 until birth (P0). Gestational absolute water intake was not different across groups (Two-Way ANOVA: *Treatment Effect* $F_{(1,53)}=0.04$, *ns*; *Exposure Effect* $F_{(2,53)}=0.08$, *ns*; *Treatment x Exposure* $F_{(2,53)}=2.70$, *ns*). No difference was seen between PTU and VEH/CON groups (Independent *t*-test: t=1.84, *ns*). Gestational relative water intake, defined as absolute water intake per number of pups per dam, differed by T4 treatment (Two-Way ANOVA: *Treatment Effect* $F_{(1,52)}=8.94$, $p=.004$), with H-DE-71-T4 dams showing significantly greater water intake compared to VEH/CON ($p=.025$). However, relative gestational water intake did not differ by exposure or interaction (*Exposure Effect* $F_{(2,52)}=1.36$, ns; *Treatment x Exposure Interaction* $F_{(2,52)}=0.89$, ns). No difference was seen between PTU and VEH/CON groups (Independent *t*-test: t=0.35, *ns*).

Similarly, there were no group differences in postpartum absolute water intake from P1-10 (Two-Way ANOVA: *Treatment Effect* $F_{(1,48)}=2.39$, *ns*; *Exposure Effect* $F_{(2,48)}=2.60$, *ns*; *Treatment x Exposure Interaction* $F_{(2,48)}=0.307$, *ns*). Absolute water intake was higher in VEH/CON compared to PTU (Independent *t*-test: t=2.53, *p*=.023). Postpartum relative water intake differed by T4 treatment (Two-Way ANOVA: *Treatment Effect* $F_{(1,48)}$=4.06, *p*=.050, but no differences were seen on post hoc analysis. Similarly, relative water intake did not differ by exposure or interaction (*Exposure Effect* $F_{(2,48)}$=0.166, ns; *Treatment x Exposure Interaction* $F_{(2,48)}$=0.05, *ns*). Relative water intake was higher in VEH/CON compared to PTU (Independent *t*-test: t=2.23, *p*=.042)(**Supplementary Table 1**).

Daily gestational weight gain was also monitored from GD7 to birth (P0). A difference in gestational absolute weight gain was observed based on the interaction of treatment and exposure (*Treatment x Exposure Interaction* $F_{(2,64)}$=3.99, *p*=.023). However, no differences were observed on Holm-Sidak post-hoc test.No differences were seen across treatment or exposure effects (Two-Way ANOVA; *Treatment Effect* $F_{(1,64)}$=0.66, *ns*; *Exposure Effect* $F_{(2,64)}$=2.41, *ns*). No difference was seen in gestational absolute weight gain between PTU and VEH/CON groups (Independent *t*-test: t=0.24, *ns*). Similarly, gestational relative weight gain differed across treatment (*Treatment Effect* $F_{(1,64)}$=5.06, *p*=.028) and exposure effect (*Exposure Effect* $F_{(2,64)}$=5.36, *p*=.007 with H-DE-71+T4 showing greater gestational relative weight gain compared to VEH/CON (*p*=.003), and L-DE-71+T4 dams (*p*=.013). No interaction effects were seen for relative weight gain (*Treatment x Exposure Interaction* $F_{(2,64)}$=1.39, *ns*). No difference was seen in gestational relative weight gain between PTU and VEH/CON groups (Independent *t*-test: t=0.50, *ns*). Percent change in absolute weight gain was not different across groups (*Treatment Effect* $F_{(1,64)}$=0.44, *ns*; *Exposure Effect* $F_{(2,64)}$=0.22, *ns*; *Treatment x Exposure Interaction* $F_{(2,64)}$=0.58, *ns*). No difference was seen in percent change in absolute weight gain between PTU and VEH/CON groups (Independent *t*-test: t=1.60, *ns*). Similarly, percent change in relative weight gain was not different across groups (*Treatment Effect* $F_{(1,63)}$=0.48, *ns*; *Exposure Effect* $F_{(2,63)}$=2.05, *ns*; *Treatment x Exposure Interaction* $F_{(2,63)}$=0.01, *ns*). No difference was seen in percent change in relative weight gain between PTU and VEH/CON groups (Independent *t*-test: t=1.94, *ns*).

Postpartum absolute weight gain from P1-10 differed across exposure effect (*Exposure Effect* $F_{(2,29)}$=6.98, *p*=.003), with H-DE-71 dams showing greater weight gain compared to VEH/CON+T4 (*p*=.038). However, no differences were observed across treatment or interaction (*Treatment Effect* $F_{(1,29)}$=3.66, *ns*; *Treatment x Exposure Interaction* $F_{(2,29)}$=0.28, ns). No difference was seen in postpartum absolute weight gain between PTU and VEH/CON groups (Independent *t*-test: t=1.73, *ns*). Similarly, postpartum relative weight gain differed across exposure effect (*Exposure Effect* $F_{(2,29)}$=5.27, *p*=.011), but post-hoc analysis showed no differences across groups. Postpartum Relative weight gain did not differ by treatment or interaction (*Treatment Effect* $F_{(1,29)}$=1.72, *ns*; *Treatment x Exposure Interaction* $F_{(2,29)}$=0.39, ns). No difference was seen in postpartum relative weight gain between PTU and VEH/CON groups (Independent *t*-test: t=0.50, *ns*). Percent change in absolute weight gain was different across exposure groups (*Exposure Effect* $F_{(2,29)}$=8.06, *p*=.002), with H-DE-71 showing a greater absolute percent change compared to VEH/CON (*p*=.016) and VEH/CON+T4 (*p*=.027). However, percent change in absolute weight gain did not differ by treatment or interaction (*Treatment Effect* $F_{(1,29)}$=3.66, ns; *Treatment x Exposure Interaction* $F_{(2,29)}$=0.51, *ns*). No difference was seen in percent change in absolute weight gain between PTU and VEH/CON groups (Independent *t*-test: t=1.66, *ns*). Percent change in relative weight gain differed by exposure effect (*Exposure Effect* $F_{(2,29)}$=5.77, *p*=.008), but no differences were seen on post-hoc analysis. Percent change in relative weight gain did not differ based on treatment or interaction (*Treatment Effect* $F_{(1,29)}$=2.01, *ns*; *Treatment x Exposure Interaction* $F_{(2,29)}$=0.230, *ns*). No difference was seen in percent change in relative weight gain between PTU and VEH/CON groups (Independent *t*-test: t=1.64, *ns*).

There were no exposure or treatment effects on secondary sex ratio or the average number of pups per litter.

**Supplementary Figure 1**

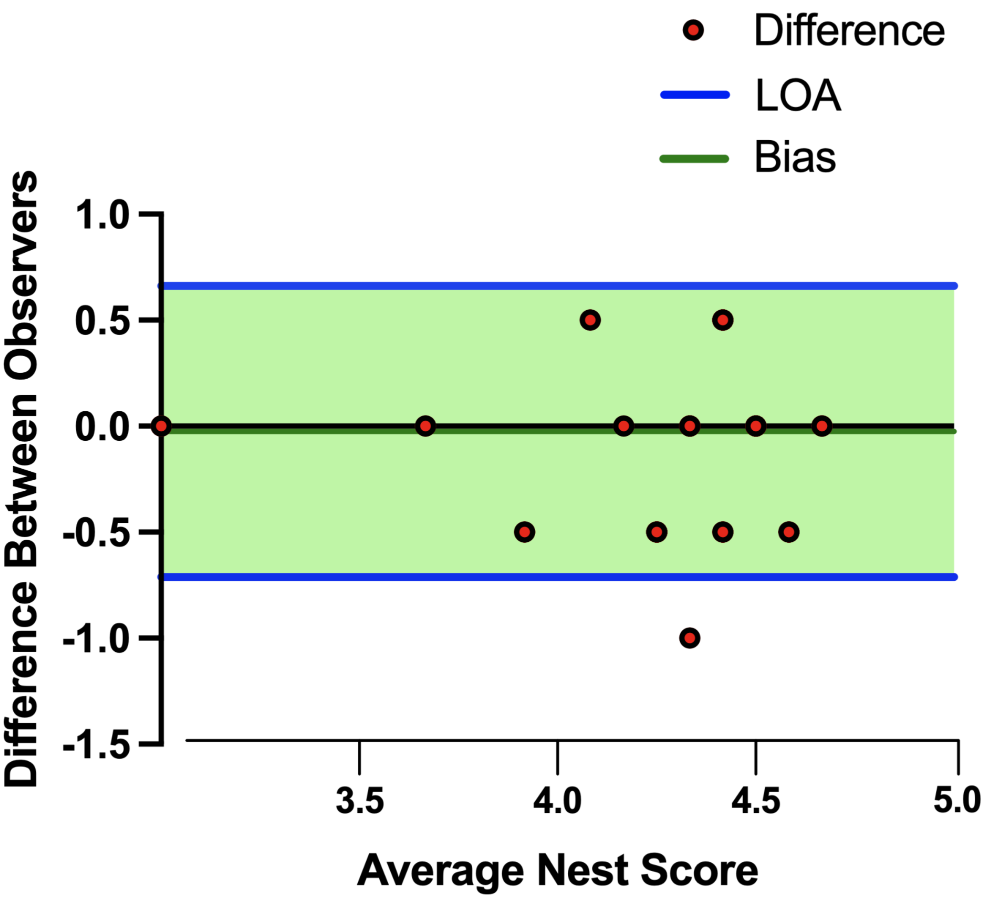


**Supplementary Figure 1. Bland-Altman plot for nest scores (Related to Figure 1).** Bland-Altman bias plot (mean±s.d.) was used to test the validity and reproducibility between two independent observers blind to exposure group. Analysis revealed a very small mean of the differences between observer scores (Bias, -0.023±0.35) and a precision measured as limits of agreement (LOA), average difference ± 1.96 standard deviation of the difference, of -0.71 to 0.66, indicating negligible skewing by either observer.

**Supplementary Table 2. Elution parameters for the analysis of THs (Related to Figure 2).**

| TIME (min) | PUMB B (%) |
|---|---|
| 0.1 | 5 |
| 3 | 5 |
| 4 | 30 |
| 5.5 | 35 |
| 6.5 | 35 |
| 7.0 | 40 |
| 8.0 | 100 |
| 10.0 | 100 |
| 10.2 | 5 |
| 13.2 | 5 |

**Supplementary Table 3. Optimized MS/MS parameters for THs (Related to Figure 2).** For each compound, ion-transitions are shown as m/z for the parent ion and two product ions (for quantification (q) and confirmation (c)). Compound optimized values for retention time ($t_R$), declustering potential (DP), collision energy (CE), and collision cell exit potential (CXP).

| Compound | $t_R$ (min)[a] | Parent ion (m/z) | Product ions (m/z) | DP (V) | CE (V) | CXP (V) |
|---|---|---|---|---|---|---|
| Target compounds | | | | | | |
| T4 | 6.90 | 778 | 732(q) | 75 | 40 | 20 |
| | | | 634 (c) | 75 | 40 | 20 |
| rT3 | 6.35 | 652 | 606 (q) | 70 | 35 | 15 |
| | | | 508 (c) | 70 | 35 | 15 |
| T3 | 6.07 | 652 | 606 (q) | 70 | 35 | 15 |
| | | | 508 (c) | 70 | 35 | 15 |
| 3,3’-T2 | 5.67 | 526 | 480 (q) | 65 | 40 | 20 |
| | | | 382 (c) | 65 | 40 | 22 |
| 3,5-T2 | 5.34 | 526 | 480 (q) | 65 | 35 | 20 |
| | | | 382 (c) | 65 | 35 | 22 |
| T1 | 5.11 | 400 | 256 (q) | 70 | 35 | 20 |
| | | | 354 (c) | 70 | 35 | 20 |
| $T_1$AM | 5.22 | 356 | 339 (q) | 75 | 40 | 35 |
| | | | 212 (c) | 75 | 40 | 25 |
| Internal standards | | | | | | |
| ML-T4 | 6.90 | 784 | 738 | 75 | 35 | 20 |
| ML-rT3 | 6.35 | 658 | 612 | 70 | 35 | 15 |
| ML-T3 | 5.67 | 658 | 612 | 70 | 35 | 15 |
| ML-3,3’-T2 | 5.11 | 532 | 486 | 65 | 40 | 25 |
| ML-$T_1$AM | 5.22 | 362 | 345 | 75 | 40 | 35 |

[a] Retention time for the Shimadzu Nexera X2 LC.

**Supplementary Figure 2**

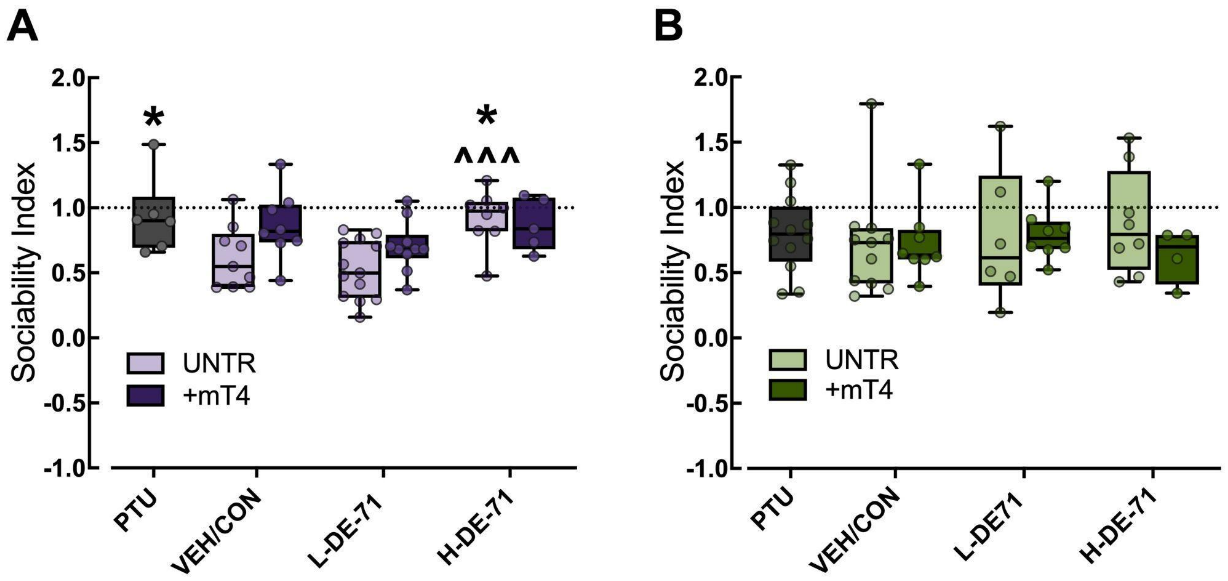


**Supplementary Figure 2. Sociability Index (Related to Figure 4).** (**A**) Sociability index in female offspring. (**B**) The sociability index in male offspring. *indicates a statistical difference vs VEH/CON in A (*$p$<.05). ^significant difference vs L-DE-71 in A (^^^$p$<.001). Values represent the means±s.e.m. $n$, 6-13/group (A) and 6-12/group (B). +mT4: maternal T4. UNTR: untreated

**Supplementary Figure 3**

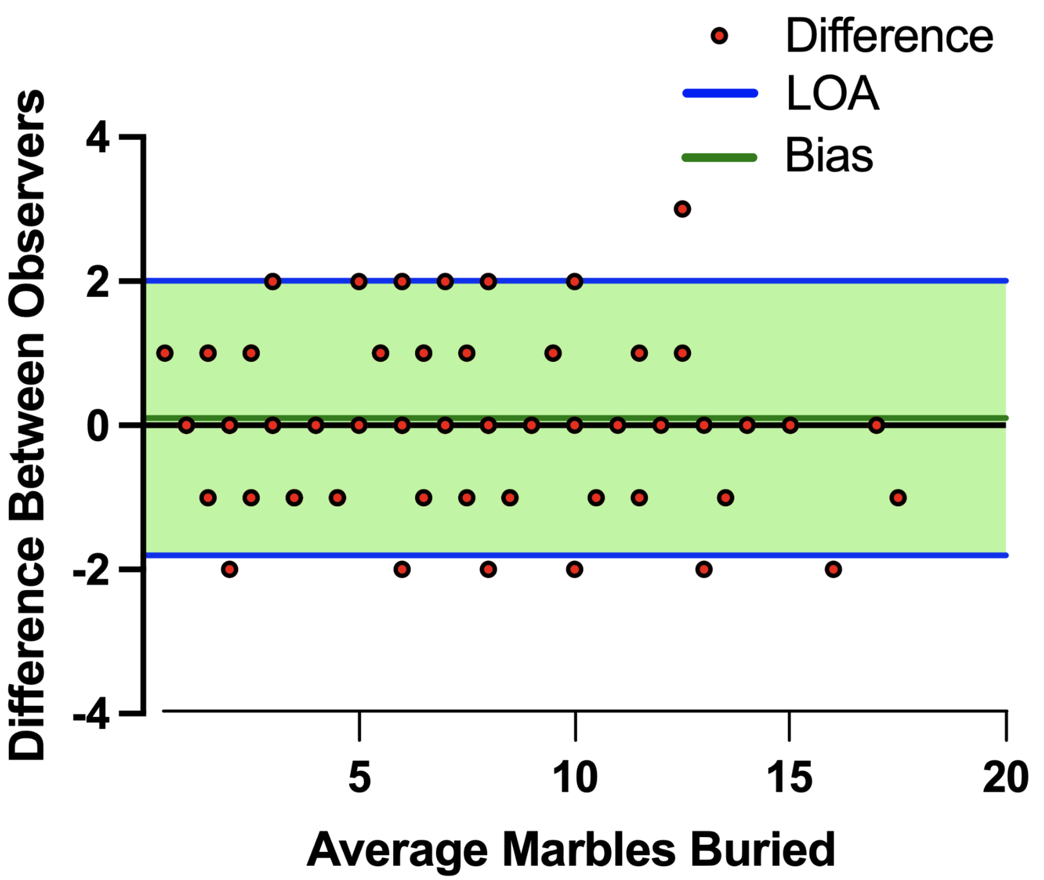


**Supplementary Figure 3. Bland-Altman plot for marble (Related to Figure 4).** Bland-Altman bias plot (mean±s.d.) was used to test the validity and reproducibility between two independent observers blind to exposure group. Analysis revealed a very small mean of the differences between observer scores (Bias, 0.10±0.97) and a precision measured as limits of agreement (LOA), average difference ± 1.96 standard deviation of the difference, of -1.8-2.2, indicating negligible skewing by either observer.

**Supplementary Table 4: RT-PCR primer sequences and efficiencies and PCR products. (Related to Figure 5)**

| Target Gene | Gene Symbol | GenBank Accession Number | Primer Sequence Forward 5’-3’ Reverse 5’-3’ | Exon target (Fwd/Rv) | E (%) | Tm°C (Fwd/Rv) | Product length (bp) | Anneal temp (°C) |
|---|---|---|---|---|---|---|---|---|
| Arginine vasopressin[a] | *Avp* | NM_009732.2 | CTC AAC ACT ACG CTC TCC GC<br>CAG CAG ATG CTT GGT CCG A | 1/1–2 | 98 | 60.8/57.9 | 173 | 55 |
| Beta actin[a] | *Actb* | NM_007393 | GATTACTGCTCTGGCTCCTAG<br>GACTCATCGTACTCCTGCTTG | 5/6 | 99.6 | 55.0/54.4 | 147 | 60 |
| Cyclic ADP ribose hydrolase | *Cd38* | NM_007646 | TGC TCC TTT GAA TGT ACT CAG T<br>AGC ACT TTT CTG ACA TCT TCC T | 1-2 | 100.9 | 57.4/57.6 | 110 | 60 |
| Oxytocin[a] | *Oxt* | NM_011025.4 | CCGAAGCAGCGTCTTTT<br>CTTGGCTTACTGGCTCTGAC | 1/2 | 96.9 | 55.7/55.5 | 131 | 60 |
| Thyrotropin-releasing hormone[b] | *Trh* | NM_009426.3 | GATTCTGGAGCCTTGCAGAC<br>GGGGATACCAGTTAGCACGA | 1/2 | 101.7 | 58.6/58.9 | 156 | 60 |

Abbreviations: *E* primer efficiency, *Tm* melting temperature, *bp* base pair

Primer sequences were obtained from: a: (Kozlova et al., 2022); b: (Swarnalatha & Abraham, n.d.)

**Supplementary Figure 4**

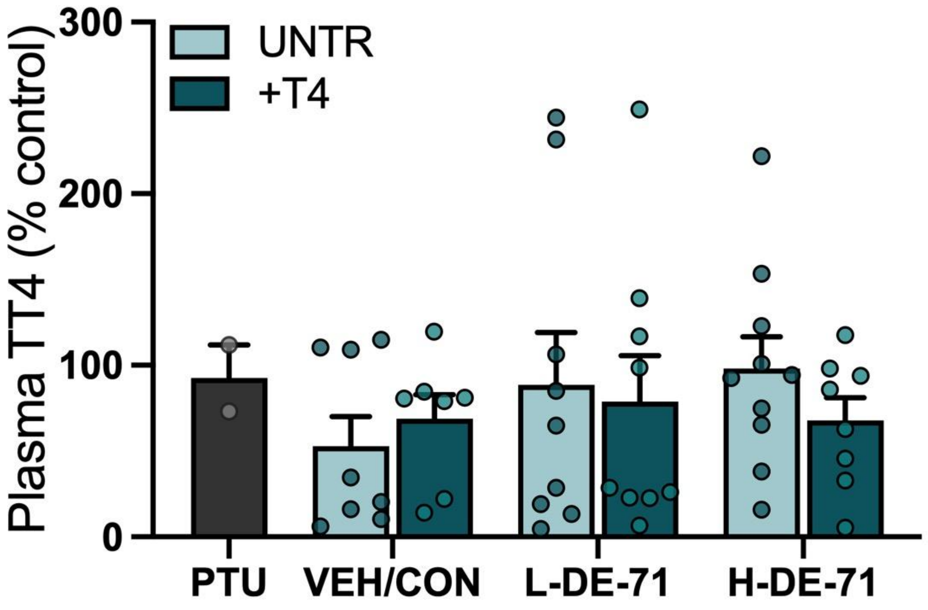


**Supplementary Figure 4. (Related to Fig 1)** Plasma total T4 in dams at P22. No statistical significance between groups and treatment. *n*, 2-10/group. UNTR: untreated

**Supplementary Figure 5**

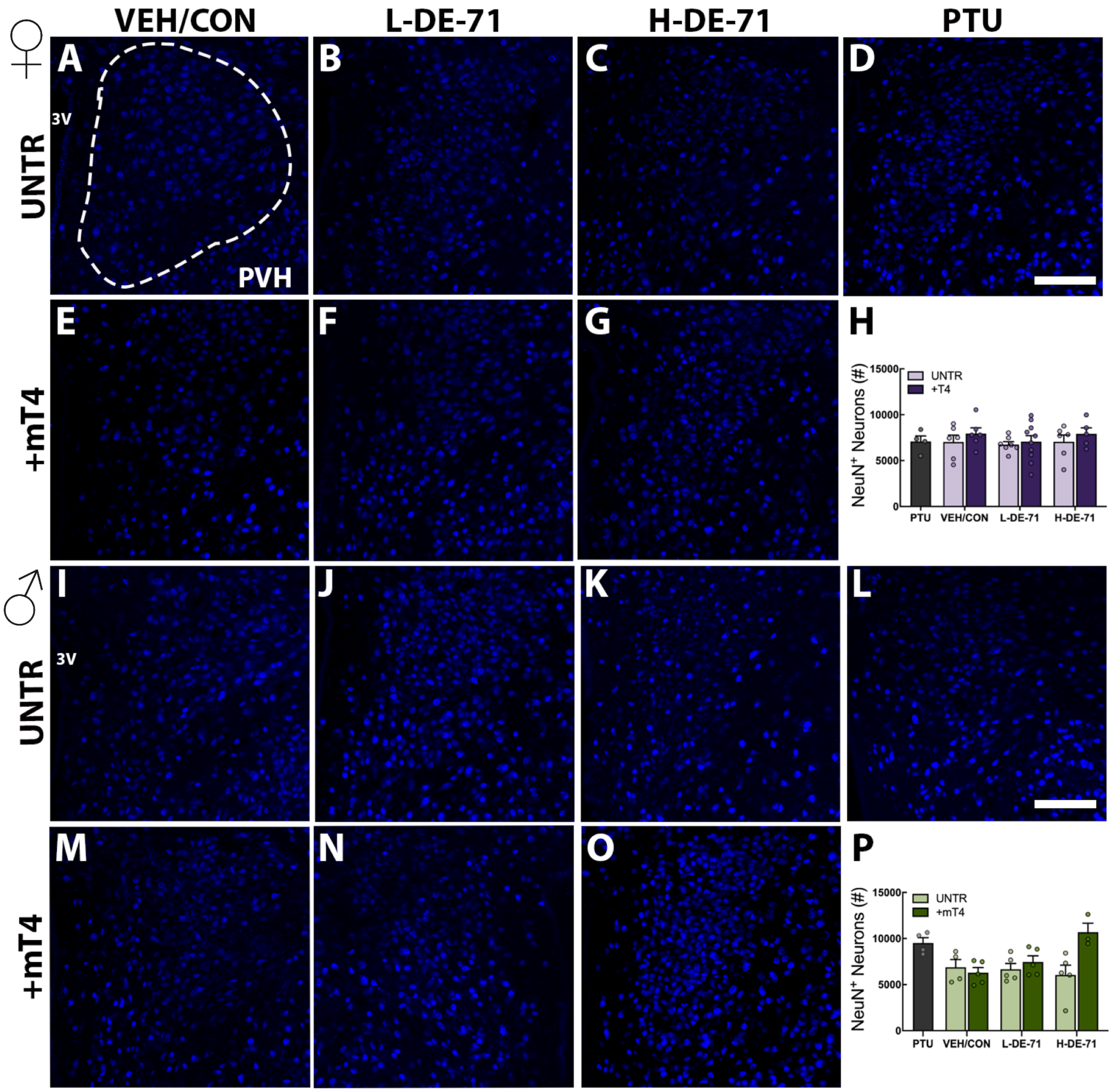


**Supplementary Figure 5 (Related to Figure 6).** NeuN+ neurons in PVH in females **(A-G)** and males **(I-O)**. Stereological quantification of Neun+ PVH neurons in females **(H)** and males **(P)**. The reduced OXT content in the PVH caused by L-DE-71 is not due to a reduction in global neuron number. *n*, 4-19/group (I); 4-10/group (H); 3-5/group (P) Scale bar, 100 microns

**Supplementary Figure 6**

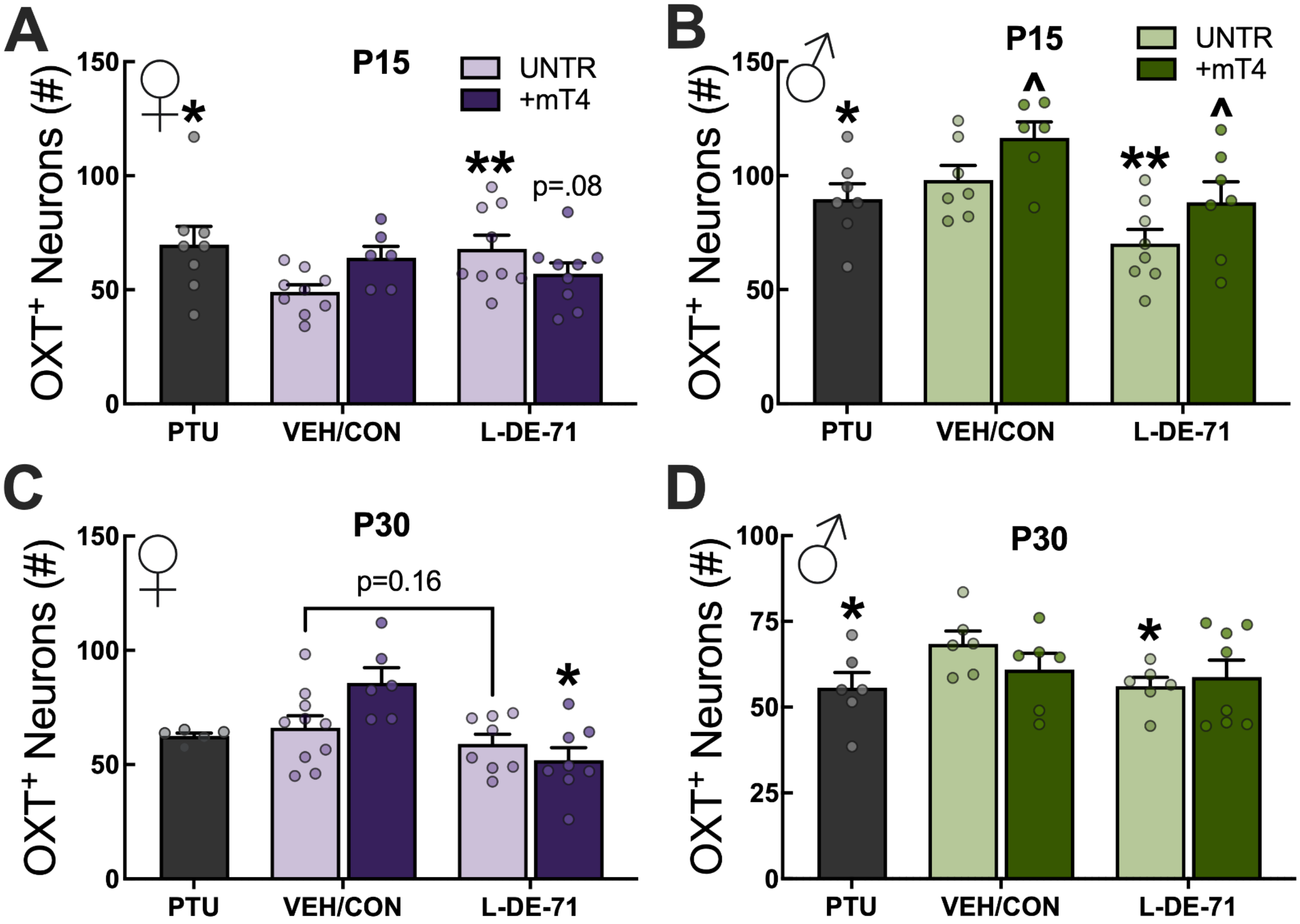


**Supplementary Figure 6 (Related to Figures 8, 9).** OXT+ neuron counts in PVH using HALO-AI digital image analysis in females at P15 (**A**), males at P15 (**B**), females at P30 (**C**), males at P30 (**D**). *n*, 6-9/group (A); 5-8/group (B); 5-10/group (C); 5-8/group (D)
